# *In situ* growth of anammox bacteria in subseafloor sediments

**DOI:** 10.1101/729350

**Authors:** Rui Zhao, José M. Mogollón, Sophie S. Abby, Christa Schleper, Jennifer F. Biddle, Desiree Roerdink, Ingunn H. Thorseth, Steffen L. Jørgensen

**Author notes:** University of Grenoble Alpes, CNRS, Grenoble INP, TIMC-IMAG, Grenoble, France. Correspondence: Rui Zhao, School of Marine Science and Policy, University of Delaware, Lewes, Delaware, USA.

## Abstract

The deep biosphere buried in marine sediments was estimated to host an equal number of microbes as found in the above oceans ^1^. It has been debated if these cells are alive and active ^2^, and their per cell energy availability does not seem to allow for net population growth ^3^. Here, we report the growth of anammox bacteria in ∼80,000 year old subsurface sediments indicated by their four orders of magnitude abundance increase in the nitrate-ammonia transition zone (NATZ). Their growth coincides with a local increase in anammox power supply. The genome of the dominant anammox bacterium from the NATZ was reconstructed and showed an increased index of replication confirming *in situ* active growth. The genome belongs to a new *Scalindua* species so far exclusively found in marine environments, which has the genetic capacity of urea and cyanate utilization and is enriched in genes allowing it to cope with external environmental stressors, such as energy limitation. Our results suggest that specific microbial groups are not only able to survive over geological timescales, but also thrive in the deep subsurface when encountering favorable conditions.

## Main text

The global cell numbers of microbes in marine sediments is estimated to be on the order of 2.9-5.4×10^29^ equaling up to 1/3^rd^ of the total prokaryotic biomass on Earth ^1,4^. A considerable portion of these cells reside beyond the bioturbation zone and constitute the marine deep biosphere ^5^. Microbial cells in the subseafloor sediments are sealed off from recruitment of new cells and fresh substrates from the surface, and therefore are thought to suffer severe energy limitations ^3^. Despite this, several lines of circumstantial evidence indicate that the deep microbial biosphere is alive ^6,7^, but with extremely slow metabolic rates ^8,9^ and long turnover times of hundreds to thousands of years ^2,10^. Although microbial growth (net biomass production) was frequently assumed ^10,11^ and recently observed in laboratory incubations ^12^, concrete evidence of *in situ* microbial growth in the marine deep biosphere is lacking.

Energy availability is considered one of the most fundamental factors limiting life, but has not been explicitly demonstrated to control the changes of microbial communities in the deep biosphere ^13^. Whereas the deep sedimentary realm is a stable environment with low energy availability, geochemical transition zones such as the sulfate-methane transition zones^14^ and oxic-anoxic transition zones^15^ are known to harbor higher microbial abundances than adjacent depths. Higher energy/power availability provided by the intensified redox reactions was invoked to explain this phenomenon. Whether this theory can be generalized to other geochemical transition zones, however, is still unknown. Here we present the geochemistry, microbial ecology, energetics, and genomic characterization of a novel *Scalindua* anammox bacterium from a nitrate-ammonia transition zone (NATZ, the sediment interval where NO_3_^-^ and NH_4_^+^ co-exist), providing compelling evidence for *in situ* microbial growth associated with increased power availabilities in ∼ 80,000 year old subsurface sediments.

We retrieved four sediment cores (2.0-3.6 meters long) from the seabed of the Arctic Mid-Ocean Ridge (AMOR) at water depths of 1653 – 3007 m (Fig. 1a and Table S1), to perform high vertical resolution geochemical measurements and microbiological analyses. All four cores exhibited similar geochemical profiles (Fig. 2a-c), summarized as follows: 1) O_2_ monotonically decreased and was depleted at a depth of 0.4-1.2 mbsf, while dissolved Mn^2+^ built up right below the oxygen depletion zone; 2) NO_3_^-^ was abundant in the oxic zone and depleted in layers below oxygen depletion depth; and 3) NH_4_^+^ was abundant in the deep anoxic sediment but undetectable in sediment above the nitrate depletion depth. Such geochemical profiles clearly indicate that each core harbors a NATZ, where both nitrate diffusing downward from the oxic zone and ammonium diffusing upward from deeper parts of the sediments are co-consumed, presumably via the anammox process (Fig. 2 and Table S1). Flux calculation suggested that most of the NH_4_^+^ flux (60-100%) diffusing from the deep anoxic sediments was consumed in the NATZ (Table S1). By revisiting meta-data of earlier studies (see Supplement Information for details), we noted that NATZ exits in many locations with water depth of 1000 to 3500 meters (Fig. 1b), and is likely widespread in the vast, yet discretely sampled, deep sea sediments.

**Figure 1.**
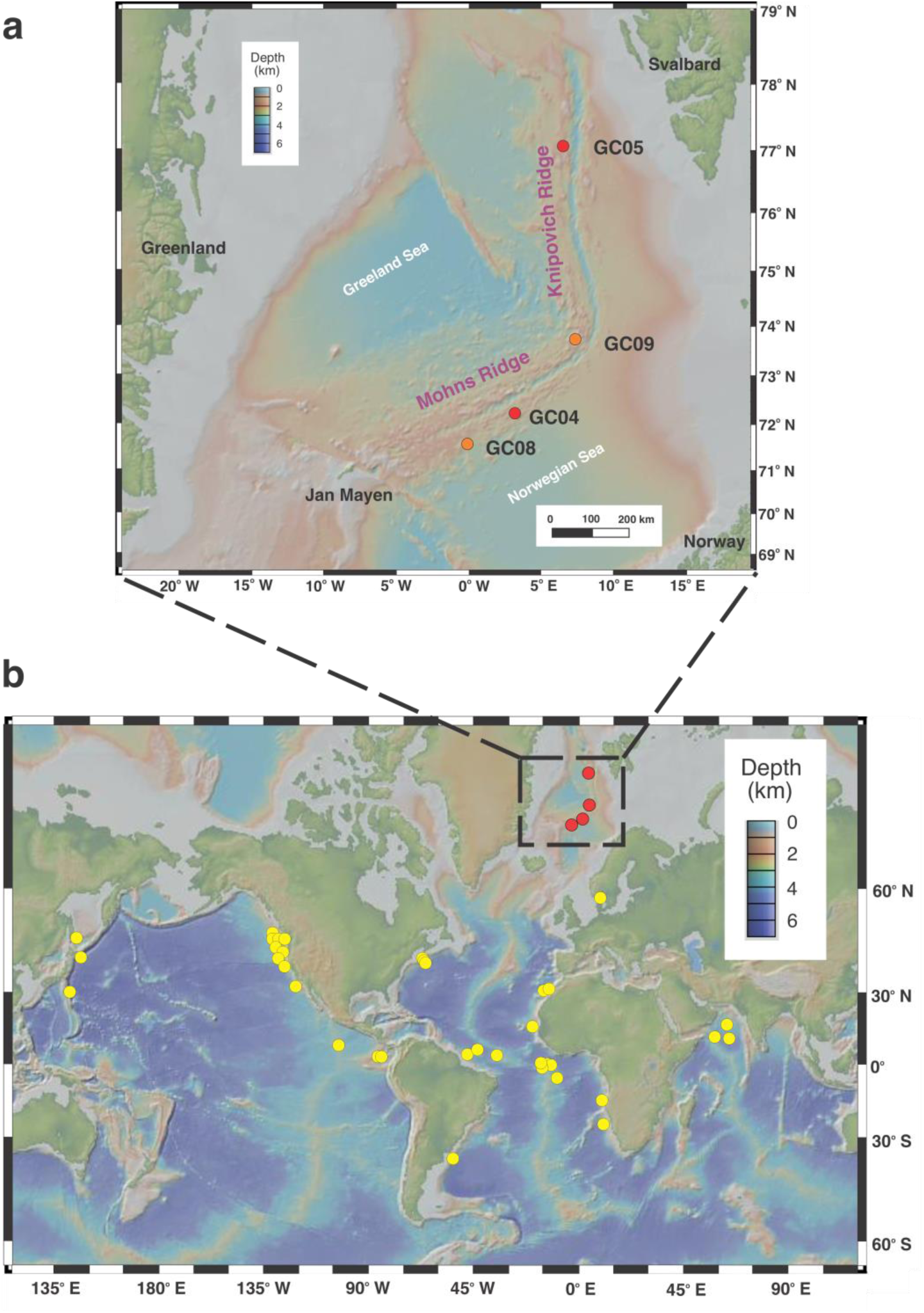
Sampling site of this study (a) and the global occurrence of nitrate-ammonium transition zone (NATZ) (b). **(a)** Bathymetry map of the Arctic Mid-Ocean Ridge highlighting Mohns Ridge and Knipovich Ridge in the Norwegian-Greenland Sea. Cores GC08 and GC09 (orange) were sampled during the CGB 2014 summer cruise. Cores GC04 and GC05 (red) were sampled during the CGB 2016 summer cruise. Map was created in GeoMapApp version 3.6.10 using the default Global Multi-Resolution Topography Synthesis (Ryan et al., 2009) basemap. **(b)** Location of marine sediments bearing an observed NATZ, which was identified based on the measured profiles of nitrate and ammonium. The box corresponds to the AMOR area shown in the upper panel (a).

**Figure 2.**
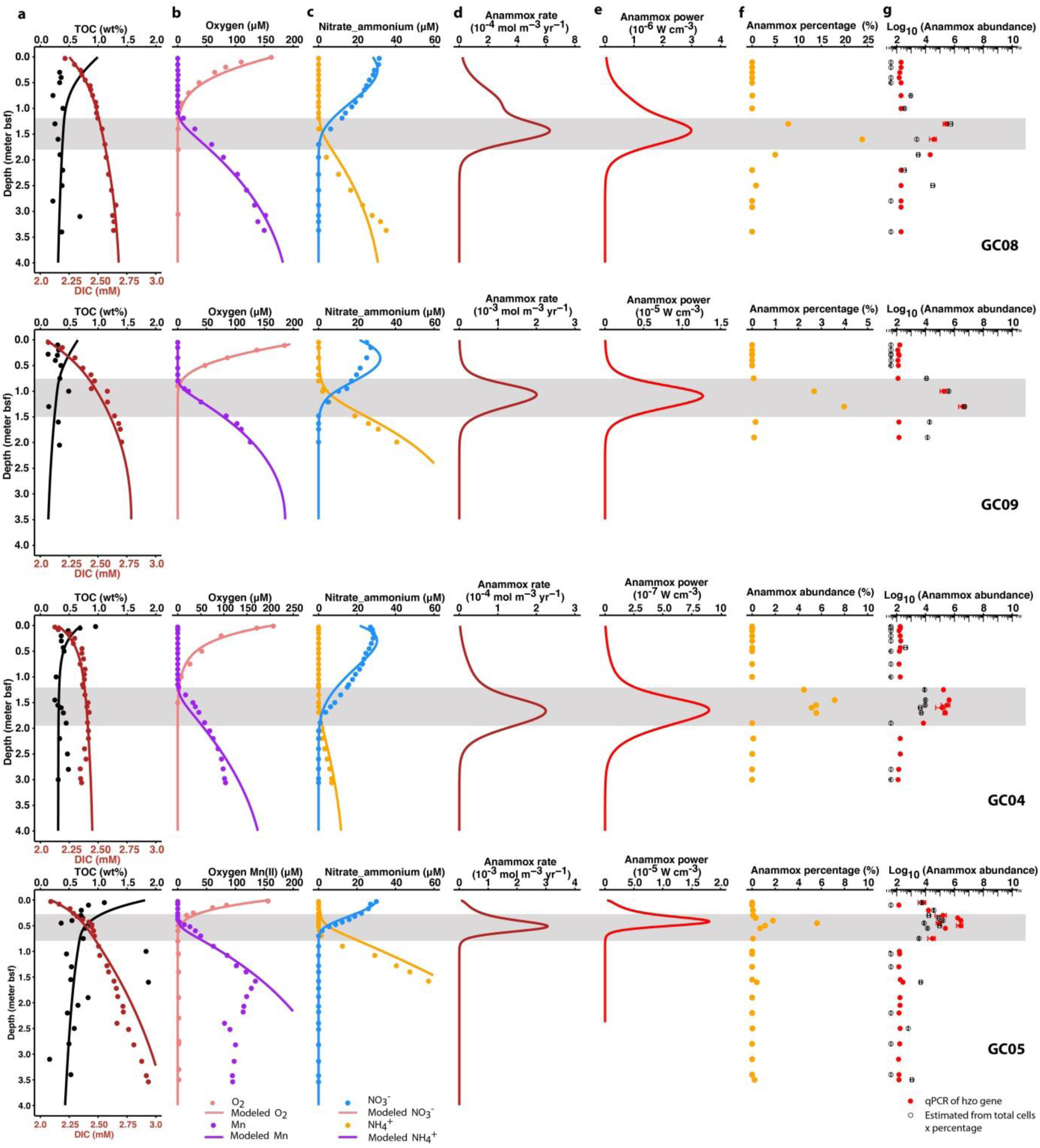
Distribution of anammox bacteria abundances and reaction rates. **a-c)** Measured (dots) and modeled (lines) depth profiles of total organic carbon (TOC), dissolved inorganic carbon (DIC), oxygen, dissolved manganese, nitrate and ammonium. **d)** Anammox rate calculated based on model simulation. **e)** Power supply of anammox calculated as the products of anammox rate and Gibbs free energy per anammox reaction presented in Supplementary Fig. S1. **f)** Percentage of anammox bacteria from the genus of *Scalindua* in the amplicon libraries. **g)** Anammox bacteria abundance quantified by two methods: 1) qPCR targeting the *hzo* gene (encoding the hydrazine dehydrogenase) (filled red dots), and 2) estimated as the products of anammox bacteria percentage in (f) and the total cell abundances quantified by 16S rRNA genes (open dots). The NATZ in each core is highlighted by a grey box.

We applied a one-dimensional reaction-transport model ^16^ to simulate the profiles and calculate the rates of various reactions including anammox. The applied boundary conditions (Table S2) and model parameters (Table S3), allowed a simulation matching the measured profiles of TOC, DIC, O_2_, NO_3_^-^, NH_4_^+^, and Mn^2+^ (Fig. 2a-c), suggesting that the modeled profiles and reaction rates provided a realistic estimation of the *in situ* geochemical processes in these cores. Although thermodynamic calculations suggest the anammox reaction to be highly favorable at most depths across all cores (Gibbs free energy up to -200 kJ mol^-1^ N; Supplementary Fig. S1), anammox is known to be inhibited in the oxic zones by oxygen and limited in the deeper, anoxic sediments due to the absence of nitrate/nitrite. In line with this, our model predicts that anammox mainly occurred in the NATZs (Fig. 2d).

16S rRNA gene sequences show that *Scalindua*, represented by three OTUs (Operational Taxonomic Units, 97% identity), was the only genus of known anammox bacteria identified in these sediments (Fig. 3a and 3c). Consistent with the predicted anammox rate, *Scalindua* accounted for up to 24% of the total community in the NATZ, but were undetectable in the upper oxic zone and below the depth of nitrate-depletion (Fig. 2f). To determine whether such relative abundance peaks in the NATZs resulted from an increase in the absolute abundance of anammox bacteria, we quantified anammox bacteria throughout the four cores by 1) quantitative PCR targeting the *hzo* gene (encoding the hydrazine oxidoreductase) and 2) calculating their abundance by multiplying the relative abundance of anammox related taxa in the total community and the total cell numbers determined by 16S rRNA gene quantification. The absolute abundances of anammox bacteria from both methods generally agreed with each other, and showed peaks (up to 3.9×10^6^ cells g^-1^) in the NATZ in all cores, consistent with the relative abundance profiles (Fig. 2g).

**Figure 3.**
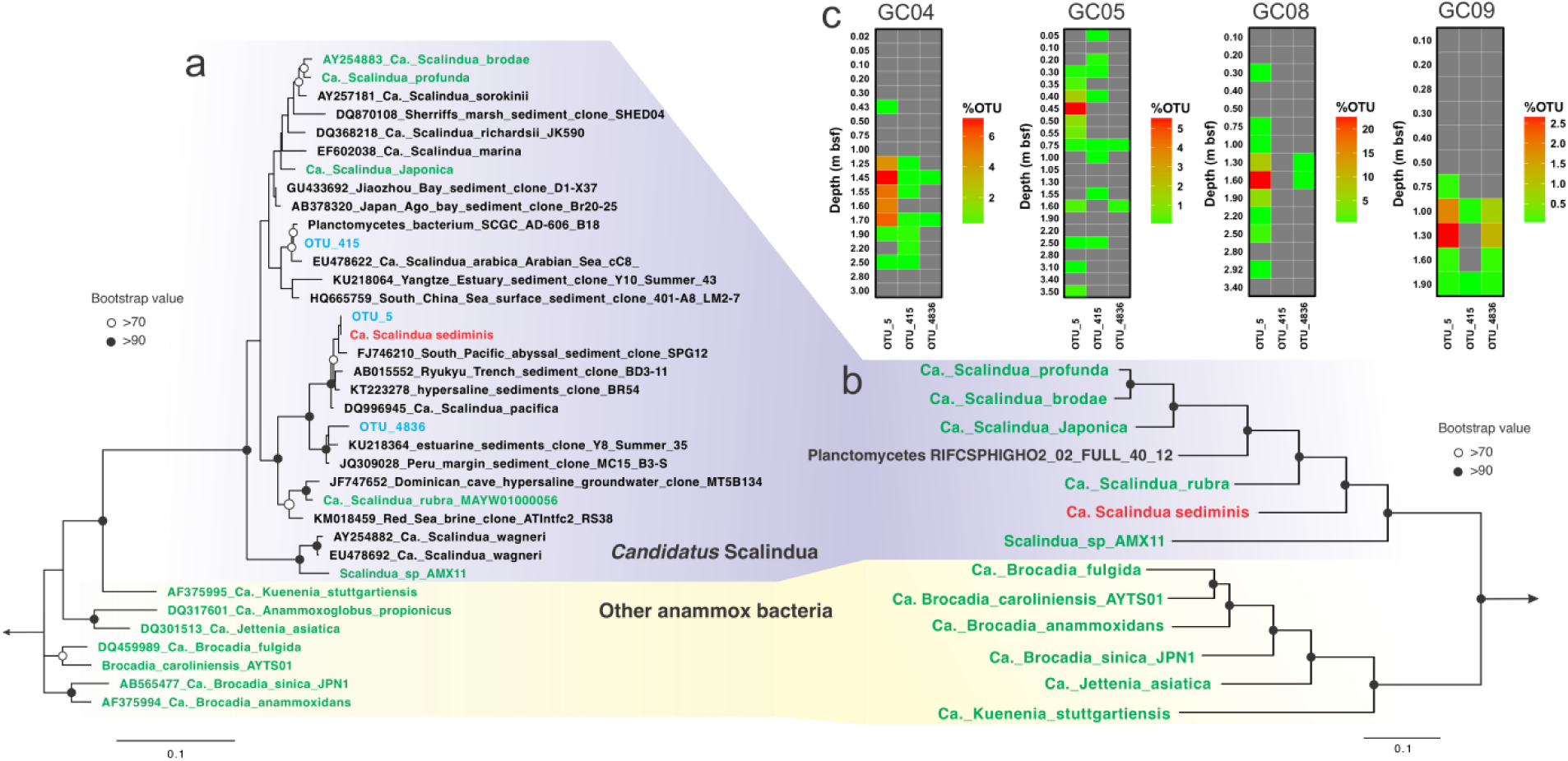
Phylogeny and vertical distribution pattern of *Scalindua* bacteria in AMOR sediments. **(a)** 16S rRNA phylogenetic tree of anammox bacteria. The three Scalindua OTUs recovered from the AMOR sediments via 16S rRNA gene amplicon sequencing are shown in blue. The tree was constructed by maximum-likelihood using RAxML with the GTAGRAMMA model. **(b)** Phylogenetic tree of ananmmox bacteria inferred from 13 concatenated ribosomal protein genes (rpL2, 3, 4, 5, 6, 14, 15, 18, 22 and rpS3, 8, 10, 17, 19). The tree was constructed by maximum-likelihood using RAxML, with PROTGAMMALG as the evolutionary model. In both (a) and (b) *Ca.* Scalindua sediminis is highlighted in red, while known anammox bacteria are highlighted in green. *Paludisphaera borealis* PX4 and *Isosphaera pallida* were used as the outgroup for both trees. Bootstrap values higher than 70 and 90 were shown on nodes with open and filled circles, respectively. The scale bars correspond to substitution per site. **(c)** Distribution of *Scalindua* OTUs in the four AMOR sediment cores. Depths of sediment horizons (meters below seafloor) are indicated on the vertical axis for each core. Note different scales are used for the anammox OTU percentages (of total community) in different cores.

To explore the factors driving the increase of anammox abundance in the subsurface, we calculated the power supply of anammox as the product of the Gibbs free energy and the modelled rate of anammox ^13^. The anammox power supply exhibited the same distribution pattern as the anammox bacterial abundance (Fig. 2e), suggesting that the increased power availability allows a higher standing stock of anammox bacteria in this zone. Given our current knowledge of the deep subsurface, this observation strongly suggests *in situ* growth. However, another scenario that could in principle explain the increases of anammox cells in the NATZs is cell migration enabled by flagellar motility (Fig. 4). To investigate this possibility further we estimated the cell-specific metabolic rates from the predicted bulk anammox rate divided by the anammox abundance. Anammox bacteria in the NATZs have cellular metabolic rates of 10^-3^-10^-1^ fmol NH_4_^+^ cell^-1^ d^-1^ (Supplementary Fig. S2), meaning that they oxidize on average 7-700 ammonium ions cell^-1^ s^-1^ (28-2800 protons cell^-1^ s^-1^). Apparently, their cellular metabolic rates are lower than the required metabolic level (10^4^-10^5^ protons per second) for the rotation of a single bacterial flagellum in *E. coli* ^17^, suggesting that these anammox bacteria do not have enough metabolic activity to fuel the flagellar rotation. Instead, we speculate the function of the flagella of anammox bacteria in the subsurface are more likely facilitating their adhesion onto particle surfaces ^18^, as the majority of microbial cells in marine sediments are particle-attached ^19^. This could provide them a protection in unfavorable conditions like the oxic zone where they could hide in anoxic microniches. Therefore, we argue based on our quantitative data that the increase of anammox bacteria abundance in the NATZs is a result of *in situ* growth rather than cell migration.

**Figure 4.**
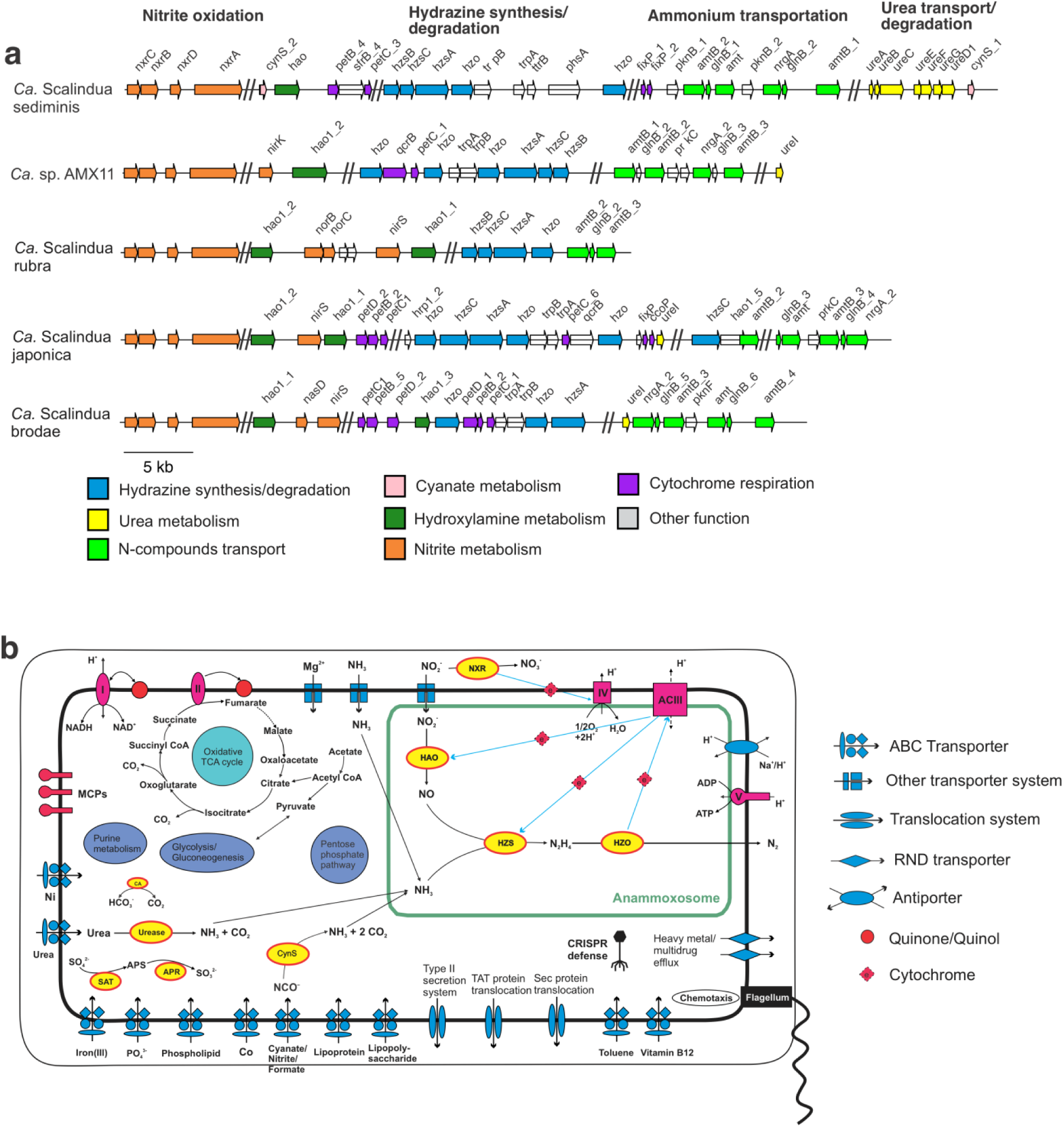
Key genome regions (a) and metabolic potential (b) of “*Canditatus* Scalindua sediminis”. **(a)** Schematic representation of key genome regions in *Ca.* Scalindua genomes. Arrows represent genes and indicate the transcriptional direction. Homologous genes are connected by lines. Genes are drawn to scale. **(b) Reconstruction of cell metabolic pathways based on the annotation of the “*Canditatus* Scalindua sediminis” genome.** Enzyme complexes of the electron transport chain are labelled with Roman numerals. The flow of electron transfer is represented by blue arrows. ACIII, alternative complex III; CA, carbonic anhydrase; CoA, coenzyme A; CRISPR, clustered regularly interspaced short palindromic repeats; SAT, sulfate adenylate transferase; APR, adenosine-5′-phosposulfate reductase; ASR, anaerobic sulfite reductase; CynS, cyanate hydratase; HZS, hydrazine synthase; HZO, hydrazine dehydrogenase; MCPs, methyl-accepting chemotaxis proteins; NIR, nitrite reductase; NXR, nitrite oxidoreductase; RND transporter, resistence-nodulation-cell division transporter; TAT, twin-arginine translation; TCA cycle, tricarboxylic acid cycle; Sec, secretion.

We performed a metagenomic analysis of the NATZ of core GC08 to study the ecophysiology and potential adaptation mechanisms of anammox bacteria in the subsurface. From the assembled and binned metagenome, we recovered a draft genome of *Scalindua* (95.5% completion). This genome was ∼3.0 Mbp, with 2,879 coding sequences across the 71 scaffolds, and thus more than 1 Mbp smaller than other known *Scalindua* genomes (Supplementary Table S4). We calculated an iRep value of 1.32 for this genome (Supplementary Table S4), suggesting that 32% of this population was in a state of active replication at the sampling time, consistent with the deduced *in situ* growth described above. The assembled (full-length) 16S rRNA gene sequence of this genome is identical to that of the dominant *Scalindua* found in our 16S rRNA amplicon analysis (Fig. 3a), suggesting that this genome represents the most dominant *Scalindua* species in the NATZ. The genome shares less than 90% 16S rRNA sequence identity and 74-81% of genomic ANI (average nucleotide identity) with previously characterized *Scalindua* species from other marine habitats, including *Ca*. S. rubra ^20^ and Ca. S. AMX11 ^21^ from seawater, *Ca.* S. japonica ^22^ and *Ca.* S. profunda ^23^ enriched from coastal sediments. Its 16S rRNA gene forms a monophyletic clade with *Ca.* S. pacifica (a genotype detected in coastal Bohai Sea sediments ^24^) and other uncultured *Scalindua* from marine sediments (Fig. 3a). Consistent with this ecotype-specific pattern, a search using the 16S rRNA gene as a query against the NCBI short reads archive (See Methods) showed that the *Scalindua* species represented by this bacterium (97% 16S rRNA nucleotide identity) were present in 120 samples (as of October 2018), all of which were marine sediments if only natural environments were considered (Supplementary Table S3). Phylogenetic analyses of concatenated ribosomal proteins (Fig. 3b) and the hydrazine synthase alpha subunit (HzsA, Supplementary Fig. S3) confirmed that this genome represents a deep-branching lineage within the genus of *Scalindua*. This genome has the complete core genetic machinery to perform anammox, including the nitrite disproportionation to nitrate by nitrite oxidoreductase (NXR) and to nitric oxide (NO) by octaheme hydroxylamine oxidoreductase (HAO) ^25^, hydrazine synthesis from NO and NH_4_^+^ catalyzed by HZS, and hydrazine degradation to N_2_ by hydrazine dehydratase (HZO) (Fig. 4). We propose a provisional taxon name for this uncultivated anammox bacterium, “*Candidatus* Scalindua sediminis”, based on its prevalence in deep marine sediments.

Notably, *Ca.* S. sediminis has the potential of utilizing urea and cyanate indicating a versatile metabolic lifestyle. For urea metabolism, it encodes a urease operon (UreABC) and a urea-specific ABC transporter, as well as several urease accessory proteins (UreDEFG) that facilitate the transportation and intracellular degradation of urea to NH_4_^+^ (Fig. 4). For cyanate metabolism, it has two copies of cyanate hydratase (encoded by *cynS*), catalyzing the degradation of cyanate to NH_4_^+^ and CO_2_ (Fig. 4 and Supplementary Fig S5). *UreC* phylogeny showed that *Ca.* S. sediminis forms a branch well-separated from known urea-utilizing nitrifiers [e.g. Thaumarchaeota, ammonia- (AOB) and nitrite-oxidizing bacteria (NOB)] (Supplementary Fig. S4), suggestinF.B. was funded in part by the WM Kecgk that *Ca.* S. sediminis had acquired the urea-utilizing capacity independently from the known urea-utilizing nitrifying organisms. Urea and cyanate are two dissolved organic nitrogen compounds ubiquitously present in seawater ^26^, and also detected in marine sediment porewater ^27^. The utilizations of these two compounds have been suggested for *Scalindua* lineages found in oxygen minimum zones based on chemical measurements ^28,29^ and supported by single-cell genome sequencing ^30^. Here we expand this observation to marine sediments, by unambiguously identifying a urease and two cyanases in a single sediment *Ca.* Scalindua genome. These two metabolic traits may not only enable *Ca.* S. sediminis to have access to alternative energy sources (i.e. urea and cyanate), but also provide it with extra ammonium to persist under the severe competition disadvantage with ammonia oxidizing Thaumarchaeota ^31^ in the upper low-oxic sediment layers.

Compared to the other five existing *Scalindua* draft genomes, *Ca.* S. sediminis is enriched in genes involved in transport and metabolism of amino acids, nucleotides, coenzymes, and lipids (Supplementary Fig. S6). In addition, *Ca.* S. sediminis encodes the lactate racemase, which is absent in other *Scalindua* genomes and could provide a rescue pathway to supply D-lactate for the cell wall synthesis during growth ^32^ when the growth-arrested status in the subsurface is relieved in the NATZ. *Ca.* S. sediminis is also unique in encoding genes for archaeal vacuolar-type H^+^-ATPase and multisubunit Na^+^/H^+^ antiporter that could decrease the energy requirement for ATP synthesis by reducing the membrane ion electrochemical potential ^33^. In addition, the RecF DNA repair pathway might also be a critical mechanism of microbial adjustment to an energy-deprived environment ^3^. Thus, the *Ca.* S. sediminis genome has extensive genetic features to invade and subsist in the energy-limiting subseafloor biosphere.

In summary we show that *Scalindua* can grow *in situ* in the subsurface NATZ, an important yet overlooked GTZ. Growth was qualitatively linked to the increased availability of power, the ultimate control of all life forms. Considering the widespread occurrence of NATZ (Fig. 1b) and other transition zones ^34^, net growth of anammox bacteria but also other organisms in the marine deep biosphere is expected to occur ubiquitously. The predominant *Ca.* S. sediminis in NATZ has genomic features that enable it to have access to alternative energy sources (e.g. urea and cyanate) and adapt to energy-limiting conditions. Our study provides evidence that certain microbial groups can maintain the dividing capacity despite being buried in the sediments for up to 80,000 years.

## Acknowledgments

We thank the chief scientist Rolf Birger Pedersen and the crew of R/V *G.O. Sars* for the retrieval of sediment cores, Anita-Elin Fedøy for the amplicon preparation, Michael Melcher, Steffen Lydvo and Gustavo Ramirez for sampling collection and DNA extraction, Jan-Kristoffer Landro for sediment carbon and nitrogen contents measurements, and Thomas Pollak for metagenome library preparation. Computational resources were made possible through the BIOMIX compute cluster (Delaware INBRE grant NIGMS P20GM103446). This work was funded by the Research Council of Norway through the Centre for Excellence in Geobiology, the K.G. Jebsen Foundation and Trond Mohns Science Foundation (to S.L.J.). S.S.A. and C.S. were supported by the Austrian Science Fund through grant P27017. R.Z. and J.F.B. was funded in part by the WM Keck Foundation.

## Data availability

All sequencing data used in this study are available in NCBI Short Reads Archive under the project number PRJNA529480. In particular, the raw metagenomic sequencing data are available in NCBI under the BioSample number SAMN11268106. The *Ca.* S. sediminis genome is available at NCBI under the accession number SAMN12415826.

## Author contributions

R.Z. and S.L.J. conceived the study. R.Z., D.R., I.H.T, and S.L.J. onboard the cruises and collected the samples. R.Z. screened the NATZ signature in literature. D.R. and I.H.T. performed the porewater extraction and analysis. R.Z. and J.M.M. performed the geochemical modeling. R.Z., S.S.A., C.S, and S.L.J. generated the metagenome data. R.Z. and S.S.A. performed metagenome assembly and binning. R.Z., S.L.J., J.F.B., and C.S interpreted results. R.Z. and S.L.J. wrote, and all authors edited and approved the manuscript.

## Competing interests

The authors declare no competing interests.

