## Supplementary Materials for "*In situ* growth of anammox bacteria in subseafloor sediments"

[1] K. G. Jebsen Centre for Deep-Sea Research, University of Bergen, Bergen, Norway.
[2] School of Marine Science and Policy, University of Delaware, Lewes, Delaware, USA.
[3] Institute of Environmental Sciences (CML), Leiden University, Leiden, Netherlands.
[4] Division of Archaea Biology and Ecogenomics, Department of Ecogenomics and System Biology, University of Vienna, Vienna, Austria.
[+] Present address: University of Grenoble Alpes, CNRS, Grenoble INP, TIMC-IMAG, Grenoble, France.

#### Materials and Methods

##### Study area and Sampling

Sediment cores used in this study were retrieved using a gravity corer from the seabed of the Arctic Mid-Ocean Ridge (AMOR) with water depths of 1653 – 3007 m, during the CGB Summer Cruise 2014 (GC08 and GC09) and 2016 (GC04 and GC05) onboard the Norwegian *R/V G.O. Sars*. GC04 and GC05 were collected from the middle section of the Knipvich Ridge, while GC08 (3.4-m long; 2,476 m water depth) and GC09 (2.0-m long; 1,653 m water depth) were collected from the Central and Northeastern end of the Mohns Ridge (deep basin Schultz), respectively (Figure 1, Table 1). Cores were taken from areas without known hydrothermal activity. Retrieved cores were immediately sectioned into 1.5-m-long whole round cores and split in halves upon arriving on deck. Oxygen concentration was measured immediately using a needle-type fiber-optic oxygen microsensor (optodes, PreSens, Regensburg, Germany) by inserting manually into sediments. The optode sensors were connected to a MICROX TX3 single channel fibre-optic oxygen meter, which was calibrated according to the manufacturer's protocols (PreSens, Regensberg, Germany). Pore water extractions were conducted with Rhizons samplers [1], from each of the 5 cm interval in the first half meter and 25 or 30 cm interval below that depth. Microbiology subsamples were taken simultaneously with porewater extraction, by using sterile 10 ml cut-off syringes from nearly identical depths as the porewater extraction, and immediately frozen at -80$^{o}$C for onshore-based DNA analysis.

Nutrient concentrations in porewater were measured onboard. Concentrations of ammonium ($NH_4^+$), nitrate ($NO_3^-$) and dissolved inorganic carbon (DIC) were analyzed colorimetrically by a Quaatro continuous flow analyzer (SEAL Analytical Ltd, Southampton, UK). The photometric indophenol method was used for ammonium measurement [2]. Nitrate was reduced to nitrite by a Cu-Cd reduction coil, and detected as a red complex [3]. Chloride ($Cl^-$) and sulfate ($SO_4^{-2}$) were measured by a Metrohm ion chromatography system. Porewater samples for metal concentrations (including Mn(II) and Fe(II)) were filtered (0.2 µm), acidified by ultrapure nitric acid to a final concentration of 3 vol%, and frozen in HDPE bottles at -20$^{o}$C until analysis. Metal concentrations were determined by Thermo IRIS ICP-OES (inductively coupled plasma optical emission spectrometry) at the University of Bergen, as described elsewhere [4].

Porosity was calculated as the weight loss of 1 cm$^{-3}$ sediment after drying at 95$^{o}$C for 24 hours, assuming a dry sediment density of 1.6 g cm$^{-3}$. Dried sediments were also used for total organic carbon (TOC) and nitrogen (TON) measurements on an element analyzer (Analytikjena multi EA® 4000, Jena, Germany), after inorganic carbon removal by adding 1 mL of phosphoric acid.

**Reaction-transport modeling**

We used the one-dimensional reaction transport model[5] to simulate the depth profiles of relevant solutes in porewater and organic carbon content in solid phase. In this study, the species explicitly modeled include oxygen, nitrate, ammonium, Mn(II), and dissolved inorganic carbon (DIC) in aqueous phase, and total organic carbon (TOC, expressed in weight percent wt%) and manganese oxide ($MnO_2$) in the solid phase. The model considers two sets of reactions: 1) the primary reactions involved in the organic matter degradation: aerobic degradation ($R_1$), heterotrophic denitrification ($R_2$), and $MnO_2$ reduction ($R_3$) and sulfate reduction ($R_4$); 2) and the secondary reactions including nitrification ($R_5$), Mn(II) oxidation with oxygen ($R_6$) and anammox ($R_7$). The model simulations assume that the geochemical profiles, including all implicit reactive intermediates, are near steady state.

Organic matter in the model was regarded to consist of 3 discrete components (the so-called 3-G model[6]), with the first two as the reactive ones and the third one as non-reactive. Aerobic respiration ($R_1$) was considered as the most favorable pathway of organic matter consumption, followed by heterotrophic denitrification ($R_2$), and $MnO_2$ reduction ($R_3$) and sulfate reduction ($R_4$), implemented through serial inhibition terms [5]. The secondary reactions ($R_5$-$R_7$) were represented through bimolecular kinetics. As nitrite is a highly active intermediate of multiple N cycle pathways, it was not explicitly simulated due to its rapid reactivity, the model assumes the anammox reaction to be a reaction between $NH_4^+$ and $NO_3^-$ following Mogollón, et al. [5]. The C/N stoichiometry of the degraded organic matter was taken as the TOC/TON ratio. The diffusion coefficients were calculated as a function of the temperature (1 $^{o}$C) and salinity (35 Practical Salinity Unit (PSU)) using the R package *marelac* (Soetaert et al., 2010a). As boundary conditions (Table 3), the model is constrained by fixed concentrations of $O_2$, $NH_4^+$, $NO_3^-$, DIC, Mn(II), $SO_4^{-2}$ and fixed organic matter flux at the sediment-water interface, and zero gradient conditions at the lower boundary of the sediment domain indicated in Table 4. The remaining model parameters (Supplementary Table S4) were calibrated by comparing the model simulation outputs with the measured depth profiles of $O_2$, $NH_4^+$, $NO_3^-$, DIC, Mn(II), $SO_4^{-2}$, and TOC (Figure 2).

The numerical solution for the partial differential equations was implemented in R following the approach outlined in Soetaert and Meysman [7]. In short, the spatial derivatives of the partial differential equations were expanded as a finite difference grid (200 equidistant layers over the sediment domain of 10 cm). After discretization, the resulting set of ordinary differential equations was integrated using the stiff equation solver *ode* implemented in R through the *deSolve* package [8].

**Occurrence of NATZ in global marine sediments**

Geochemical profiles indicating a nitrate-ammonium transition zone (NATZ) in marine sediments, marking the narrow overlap interval where downward diffusing nitrate encounter the upward diffusing ammonium, was previously reported in the literature [e.g. [9-22]]. Nitrate and ammonium profiles were obtained from published works using the online tool WebPlotDigitizer (http://automeris.io/WebPlotDigitizer), if they were not available in public databases. Additional unpublished sediment nitrate and ammonium profiles were obtained from the PANGAEA database (www.pangaea.de) by searching using the combination of the following key words: “marine sediment”, “ammonium”, and “nitrate”. Profiles were manually checked and those containing data that were too sparse points (<6) were discarded. All sites that harbor a clear NATZ were included in the global map prepared using GeoMapApp [23].

**Calculation of Gibbs free energy and power supply of anammox**

The standard Gibbs free energy ($\Delta G_r^0$) was calculated using the thermodynamic data of standard Gibbs free energy of formation of each reactant/product that corrected to near *in situ* pressure and temperature in the R package *CHNOSZ* [24]. Gibbs free energy of anammox was then calculated based on the modeled concentrations of $NH_4^+$ and $NO_3^-$, following the description in LaRowe and Amend [25]. $N_2$ concentration in sediment porewater was not measured, but assumed to be constant at $10^{-5}$ µM throughout the cores. The final values were expressed in the unit of kJ per mole of electron transferred, kJ (mol $e^-)^{-1}$, assuming six electrons transferred per anammox reaction.

Following the notion proposed in LaRowe and Amend [25], the power supply of anammox reaction, $P_s$, is calculated using the following equation:

$P_s = \Delta G_{r*} R$

where $\Delta G_r$ is the Gibbs free energy of anammox, $R$ is the anammox rate predicted from the reaction-transport model.

**DNA extraction**

DNA for amplicon sequencing and qPCR was extracted from ~0.5 of sediment per sample using PowerLyze DNA extraction kits (MOBIO Laboratories, Inc.) with the following minor modifications: 1) The lysis tube was replaced by G2 tubes (Amplikon, Odense, Denmark), and 2) lysis tubes were water bathed for 30 min at 60 °C prior to bead beating (at the speed 6.0 for 45 seconds) using a FastPrep-24 instrument (MP Biomedicals). A blank extraction was carried out in parallel with each extraction batch following the same procedure without sediment addition. DNA was eluted into 80 µL of molecular grade double-distilled $H_2O$ ($ddH_2O$) and stored at -20 $^oC$ until analysis. DNA for metagenomic sequencing was extracted from ~ 7 g sediment (0.7 g sediment in each of the 10 tubes per sample) following the procedure described above, except the final elution step: The DNA extracts from each sample were iteratively eluted from the 10 spin columns into 100 μL of $ddH_2O$ for further analysis.

**Quantification of total microbial community and anammox bacteria**

Abundance of anammox bacteria were quantified using qPCR by targeting the *hzo* gene (encoding the hydrazine dehydrogenase responsible for the degradation of hydrazine to $N_2$) using the primer pair hzoF1/hzoR1 [26], following the procedure described elsewhere [27]. In addition, archaeal and bacterial 16S rRNA genes were quantified as described in Jørgensen and Zhao [28]. Total cell abundance was estimated from 16S rRNA gene copies, assuming 4.6 copies of 16S rRNA genes for each bacterial genome, and 1.7 copies in each archaeal genome [29]. Following the method proposed by Props, et al. [30], anammox abundance was also calculated as the product of the total cell abundance and the percentage of the genus of *Candidatus* Scalindua in the total community assessed by amplicon sequencing (see description below).

**Amplicon sequencing and sequence analysis**

16S rRNA genes were amplified using the primer pair 515F/806R in a two-round amplicon preparation [31], with an optimal PCR cycle number in the first round to minimize over-amplification, and with the barcode attached in the second round of PCR. Amplicon libraries were sequenced on an Ion Torrent Personal Genome Machine in the Biodiversity Laboratory, University of Bergen, Norway. Sequencing reads were quality filtered and trimmed to 220 bp using the USEARCH pipeline [32] and chimera were detected and removed using UCHIME. Trimmed reads were clustered into operational taxonomy units (OTUs) at >97% nucleotide sequence identity using UPARSE [33]. OTUs found in the two blank controls (53 and 71 OTUs observed in the two controls respectively) were removed. Samples were subsampled to 20,000 reads for each sediment horizon with the –otutab-norm command in USEARCH v.10 [33]. The taxonomic classification of OTUs was performed using the lowest common ancestor algorithm implemented in the CREST package [34] with the SilvaMod128 database (September 2016 release) as reference. The relative abundance of anammox bacteria was taken as the sum of the percentages of *Scalindua* OTUs, and visualized in heatmaps generated using the R package *ggplot2* [35].

**Metagenomic sequencing and analysis**

DNA was sheared into 400 bp fragments using Covaris, and paired-end libraries were constructed using a Nextera DNA Flex Library Prep kit (Illumina). Metagenomic libraries were sequenced (2×150 bp) by an Illumina Hiseq 2500 sequencer at the Vienna Biocenter Core Facilities GmbH (Vienna, Austria). Quality of the reads and presence of adaptor sequences were checked using FastQC v.0.11.5 [36]. Then the sequencing data were processed with Trimmomatic v.0.36 [37] to trim read-through adapters (ILLUMINACLIP:TruSeq2-PE.fasta:2:30:10), trim low quality base calls at the starts and ends of reads (LEADING:3, TRAILLING:3), remove reads that had average phred score lower than 25 in a sliding window of 10 bp (SLIDINWINDOW:10:25), and finally remove reads shorter than 100 bp (MINLEN:100). The overall quality of processed reads was evaluated in a final check with FastQC v.0.11.5, to ensure only high-quality reads were used in the downstream analysis.

**Assembly and genome binning**

The quality-controlled paired-end reads were *de novo* assembled into contigs using Megahit v.1.1.2 [38] with the k-mer length varying from 27 to 117. Contigs larger than 1000 bp were into automatically binned with MaxBin2 v2.2.5 [39] using the default parameters. The quality of the obtained genome bins was assessed using the option "lineage_wf" of CheckM v.1.0.7 [40], which uses lineage-specific sets of single-copy genes (SCGs) to estimate completeness and contamination and assigns contamination to strain heterogeneity if amino acid identity is >90%. Genome bins of >50% completeness were manually refined using the gbtools [41] based on the GC content, taxonomic assignments, and differential coverages in different samples. Coverages of contigs in each sample were determined by mapping trimmed reads onto the contigs using BBMap v.37.61 [42]. Taxonomy of contigs were assigned according to the taxonomy of the single-copy marker genes in contigs identified using a script modified from blobology [43] and classified by BLASTn [44]. SSU rRNA sequences in contigs were identified using Barrnap [45], and classified using VSEARCH [46] with the SILVA 132 release [47] as the reference. To improve the quality of the genome of *Ca.* Scalindua sedimins, the metagenome reads of the samples GC08_160cm were mapped onto the contigs using BBmap [42], and the aligned reads were re-assembled using SPAdes v.3.12.0 [48]. After manual removal of contigs shorter than 1000 bp, the resulting scaffolds were visualized and re-binned using gbtools [49] as described above. The quality of the resulting *Scalindua* genome was checked using the CheckM v.1.0.7 "lineage_wf" command again, based on the Planctomycetes marker gene set (automatically selected by CheckM).

**Genome annotation**

Genes in the genome of *Ca.* Scalindua sediminis were predicted using Prodigal [50]. Genome annotation was conducted using Prokka v.1.13 [51], eggNOG [52], and BlastKoala [53] using the KEGG database. The functional assignments of genes of interest were also confirmed using BLASTp against the NCBI RefSeq database. The metabolic pathways were reconstructed using KEGG Mapper [54].

**Phylogenetic analyses**

All available high-quality anammox bacterial genomes were downloaded from NCBI and included in phylogenomic analysis. The phylogenomic analysis was based on marker genes consisting of 13 syntenic ribosomal proteins (rpL2, 3, 4, 5, 6, 14, 15, 18, 22 and rpS3, 8, 10, 17, 19) that have been demonstrated to undergo limited lateral gene transfer [55]. These selected proteins, among the conservative single-copy ribosomal proteins included in Campbell, et al. [56], were identified in Anvi'o v.5.4 [57] using Hidden Markov Model (HMM) profiles, following the procedure outlined at (http://merenlab.org/2017/06/07/phylogenomics/). Sequences were aligned individually using MUSCLE [58], and alignment gaps were removed using trimAl [59] with the mode of "automated". Individual alignments of ribosomal proteins were concatenated. The maximal likelihood phylogenetic tree was reconstructed using RAXML v.8.2.8 with the PROTGAMMALG model [60] or FastTree [61]. The trees generated using the two methods showed substantial agreement with each other.

A maximum likelihood phylogenetic tree of the 16S rRNA gene was also constructed for known anammox bacteria and close relatives of the three *Scalindua* OTUs identified via BLASTn [44] in NCBI. Sequences were aligned using MAFFT-LINSi [62] and the phylogeny was inferred using RAxML v.8.2.8 [60] with GTRGAMMA as the evolutionary model and 1000 fast bootstrap replicates. The short *Scalindua* OTUs amplicon sequences were placed into the phylogenetic tree using the Evolutionary Placement Algorithm [63] implemented in RAxML.

For the phylogeny of HzsA (hydrazine synthase subunit alpha), the genomes of known anammox bacteria were downloaded from NCBI, annotated using Prokka v1.13 [51], and the HzsA amino acid sequences were extracted. Additional HzsA sequences of uncultured anammox deposited in NCBI were also identified using BLASTp [44] using the HzsA sequence of *Ca.* S. sediminis as the query. Sequences were aligned using MAFFT-LINSi [62] and the maximum likelihood phylogenetic tree was inferred using RAxML v.8.2.8, with the GTRGAMMALG as the evolutionary model and 1,000 fast bootstrap replicates.

For the phylogeny of UreC (urease gamma subunit, also the catalytic subunit), the sequence of *Ca.* S. sediminis used as the query in the BLASTp [44] search in NCBI (>50% similarity were retained), to identify its close relatives. These sequences was aligned using MAFF-LINSi [62] with reference sequences from Koch, et al. [64], and complemented with known nitrifiers (e.g. ammonia-oxidizing bacteria (AOB) from the genera of *Nitrosospira*, *Nitrosomonas*, *Nitrososcoccus*, nitrite-oxidizing bacteria (NOB) from *Nitrospira* and *Nitrospina*, and ammonia-oxidizing archaea (AOA) from the phylum Thaumarchaeota). For the CynS encoding cyanase (i.e. cyanate dehydrogenase), the two copies CynS sequences of *Ca.* S. sediminis were aligned using MAFFT-LINSi [62] with reference sequences from Palatinszky, et al. [65] and their close relatives in GenBank identified via BLASTp with similarity threshold of 50%. Both alignments were then trimmed using trimAl [59] with the mode of “automated”. Maximum likelihood phylogenetic trees were reconstructed using with IQ-tree v.1.5.5 [66] under the LG+C20+F+G substitution model with 1,000 ultrafast bootstraps. All phylogenetic trees were visualized and branches were collapsed using FigTree (http://tree.bio.ed.ac.uk/publications/), prior to figure preparation in CorelDraw 2019.

**Comparative genomic analysis of *Scalindua***

Genomes of *Ca.* S. rubra [67], *Ca.* S. brodae [68], *Ca.* S. Japonica [69], Ca. S. AMX11 [70], and *Ca.* S. sediminis (recovered in this study) were included in the comparative genomic analysis using Anvio v.5.4 [57] according to the workflow described at (http://merenlab.org/2016/11/08/pangenomics-v2/). All genomes were annotated using Prokka v.1.13 [51] and BLASTp using the Clusters of Orthologous Groups of proteins (COG) [71] as the reference database. Comparative genomic information were visualized using the program anvi-interactive in Anvio v.5.4 [57]. The specific metabolic characteristics inferred from the annotations of genes with known homologs, and identified with the pangenomic analysis are discussed in the main text. The arrangements of functional genes encoding enzymes of key processes, including urea utilization, cyanate degradation, hydrazine synthesis, hydrazine degradation, nitrite oxidation, and ammonia transportation were visualized using the R package *genoPlotR* [72].

**iRep calculation**

The index of replication (iRep) was calculated for the bin *Ca.* Scalindua sediminis only in the NATZ of GC08 (160 cm bsf), because the coverages of this genome in other depths are too low to detect any changes. Settings and thresholds were applied as recommended [73] using Bowtie2 [74] and the iRep script (https://github.com/christophertbrown/iRep) with the default settings.

**Global distribution of *Ca.* S. sediminis-like anammox**

The occurrence of *Ca.* Scalindua sediminis-like anammox in natural environments was assessed using IMNGS [75] with the full-length 16S rRNA gene sequence as query. Reads with length longer than 200 bp and nucleotide sequence identity higher than 97% to the query were included as matching reads. Samples with matching reads less than 10 were discarded. Only natural environments with matching reads proportion higher than 0.05% were included.

#### Supplementary Figures

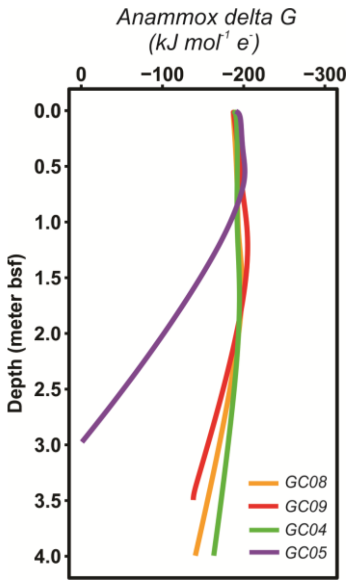

**Figure S1. Gibbs free energy of anammox reaction per mole of electron transfer.** The values were calculated based on the simulated profiles of relevant reactants and products. Positive values in the deep part of GC05 were not shown.

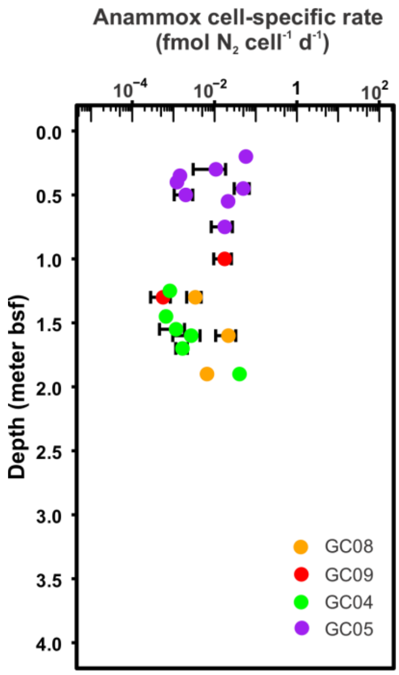

**Figure S2. Cell-specific rate of anammox bacteria in NATZ.** Cell-specific rate of anammox were calculated by dividing the modeled bulk anammox reaction rate by the anammox cell number quantified by qPCR targeting the *hzo* gene. Error bar derived from the triplicate quantification of anammox cell numbers using qPCR.

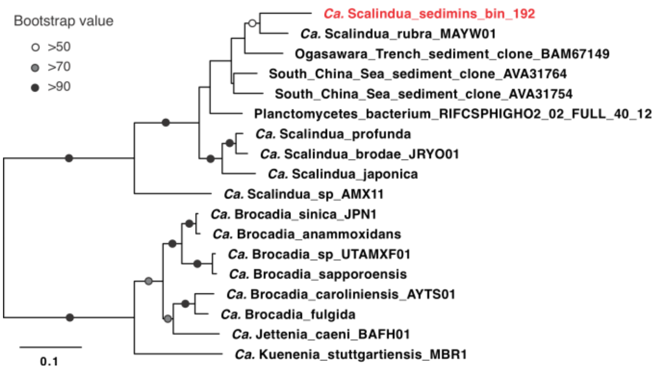

**Figure S3. Maximum-likelihood phylogenetic tree of anammox *hzsA* gene (encoding the hydrazine synthase alpha subunit).** Phylogeny was reconstructed using the PROCATJTTF model in RAxML with 1000 fast bootstraps. The *Candidatus* Scalindua sediminis genome was highlighted in red. Bootstrap values of >50 are shown with symbols listed in the legend. The scale bar shows estimated sequence substitutions per residue.

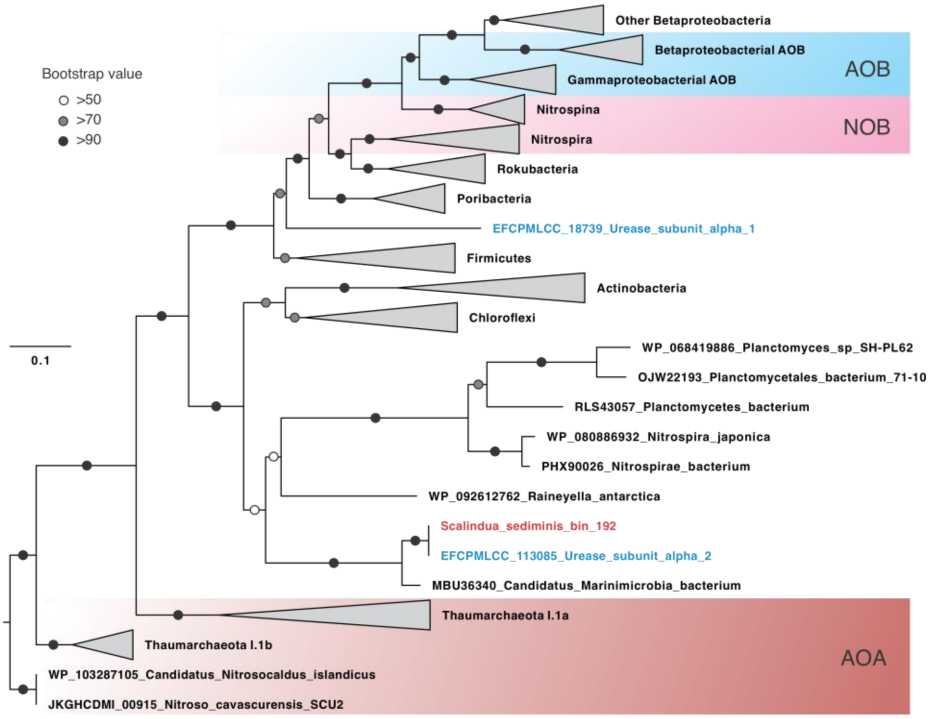

**Figure S4. Maximum-likelihood phylogenetic tree of *ureC* gene (encoding the catalytic subunit of urease)**. The phylogeny was reconstructed using IQ-tree v.1.5.5 under the LG+C20+F+G substitution model with 1,000 ultrafast bootstraps. The two *ureC* sequences detected in the metagenome assembly before binning were shown in blue and the one of *Ca.* S. sediminis are highlighted in red. Clades of nitrifying groups (i.e. AOA, AOB, and NOB) were highlighted in colored shaded boxes. Bootstrap values of >50 are shown with symbols listed in the legend. The scale bar shows estimated sequence substitutions per residue.

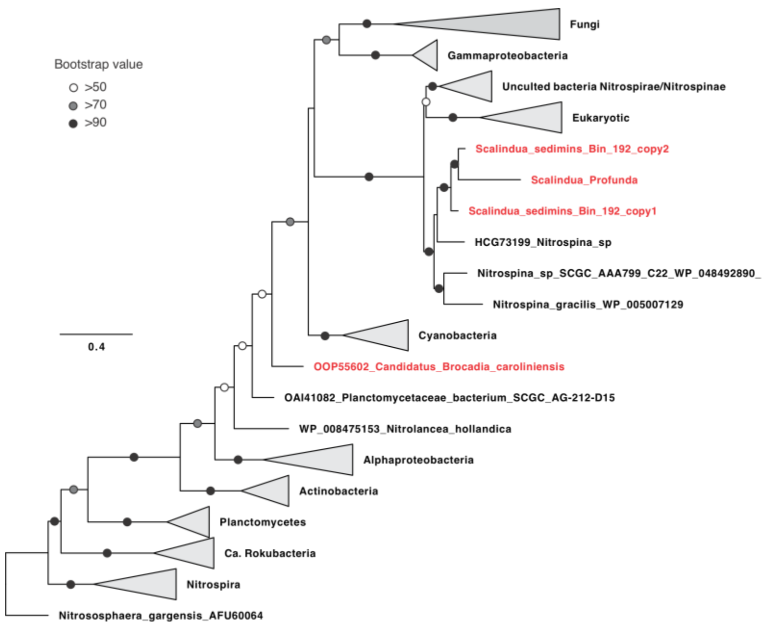

**Figure S5. Maximum-likelihood phylogenetic tree of all available full-length *cynS* gene (encoding the cyanate hydratase) amino acid sequences (~150 aa)**. Two copies of *cynS* gene detected in the *Ca*. S. sediminis genome and other anammox bacteria were highlighted in red. Phylogeny was reconstructed using IQ-tree v.1.5.5 under the LG+C20+F+G substitution model with 1,000 ultrafast bootstraps. The scale bar shows estimated sequence substitutions per residue.

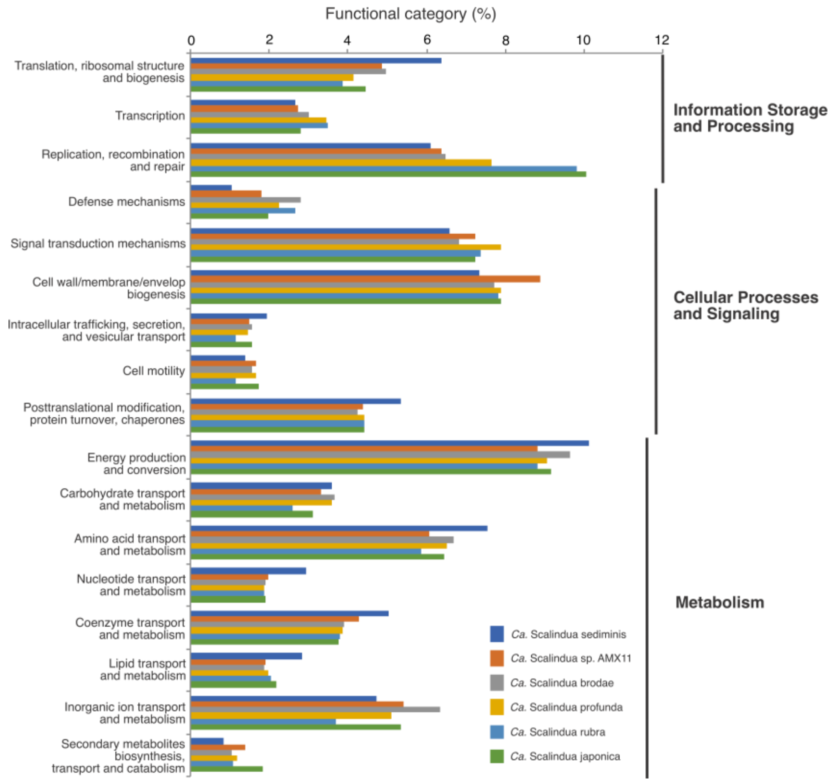

**Figure S6. Functional classification of protein-coding genes from *Ca.* Scalindua sediminis compared to other *Candidatus* Scalindua genomes.** The bar chart represents the percentage of protein-coding genes classified by eggNOG functional categories. Functional categories below 1% were excluded.

### Supplementary Tables

**Table S1. Properties of study sites considered in this study**

| Sediment core | GC08 | GC09 | GC04 | GC05 |
|---|---|---|---|---|
| Latitude (N) | $71^{o}97'$ | $73^{o}70'$ | $72^{o}16'$ | $76^{o}55'$ |
| Longitude (E) | $0^{o}10'$ | $7^{o}34'$ | $1^{o}42'$ | $7^{o}7'$ |
| Water depth (m) | 2,476 | 1,653 | 2,668 | 3,007 |
| Organic matter content (wt %) | 0.3-0.6 | 0.2-0.5 | 0.3-1.0 | 0.3-1.8 |
| Sedimentation rate (m/yr) | 2.0E-5 | 2.0E-5 | 2.0E-5 | 2.5E-5 |
| Depth of NATZ* | 1.2-1.7 | 0.8-1.5 | 1.5-2.0 | 0.5-0.7 |
| Nitrate flux to NATZ** | 0.27 | 0.24 | 0.22 | 0.64 |
| Ammonium flux to NATZ** | 0.19 | 0.40 | 0.04 | 0.82 |
| % of ammonium flux in NATZ | 100% | 60% | 100% | 78% |

*, in the unit of meters below seafloor (mbsf)

**, in the unit of mmol $m^{-2}$ $yr^{-1}$

**Table S2. Species and boundary conditions (BC) at the sediment-water interface (SWI) used in the reaction-transport model**

| Name | Symbol | BC SWI Type (Unit) | BC SWI Value | | | |
|---|---|---|---|---|---|---|
| | | | **GC08** | **GC09** | **GC04** | **GC05** |
| Total organic carbon flux | $CH_2O$ | Flux (mol $m^{-2}$ $yr^{-1}$) | 8.7E-3 | 1.42E-2 | 9.7E-3 | 2.0E-2 |
| Manganese oxide flux | $MnO_2$ | Flux (mol $m^{-2}$ $yr^{-1}$) | 6E-5 | 2.0E-5 | 4.0E-5 | 1.0E-5 |
| Oxygen | $O_2$ | Concentration (µM) | 165 | 225 | 205 | 160 |
| Ammonium | $NH_4^+$ | Concentration (µM) | 0.1 | 0.1 | 0.1 | 0.1 |
| Nitrate | $NO_3^-$ | Concentration (µM) | 25 | 21 | 21 | 30 |
| Manganese | Mn(II) | Concentration (µM) | 0.1 | 0.1 | 0.1 | 0.1 |
| Sulfate | $SO_4^{2-}$ | Concentration (µM) | 28 | 28 | 27.8 | 27.7 |
| DIC | $HCO_3^-$ | Concentration (mM) | 2.5 | 2.1 | 2.2 | 2.18 |

**Table S3. Parameter values used in the reaction-transport model**

| Name | Symbol | Unit | GC08 | GC09 | GC04 | GC05 |
|---|---|---|---|---|---|---|
| Sediment domain | L | cm | 500 | 350 | 500 | 600 |
| Solid burial velocity at compaction | ω | cm ky-1 | 2 | 5 | 2 | 2.5 |
| TOC degradation constant C1 | kfox | 1 $yr^{-1}$ | 3.0E-5 | 6.5E-5 | 6.0E-5 | 9.0E-5 |
| TOC degradation constant C2 | kfox2 | 1 $yr^{-1}$ | 1.0E-6 | 2.0E-5 | 2.0E-6 | 8.0E-6 |
| Nitrification rate constant | $k_3$ | $\mu M^{-1}$ $yr^{-1}$ | 150 | 150 | 150 | 150 |
| Mn oxidation rate constant | | $\mu M^{-1}$ $yr^{-1}$ | 110 | 110 | 110 | 110 |
| Anammox rate constant | | $\mu M^{-1}$ $yr^{-}$ | 50 | 50 | 50 | 200 |
| Bioturbation coefficient | $D_{b,0}$ | cm $yr^{-1}$ | 0 | 0 | 0 | 0 |
| Biomixing half depth | $z_{mix}$ | cm | 3 | 3 | 3 | 3 |
| Biomixing attenuation | $Z_{att}$ | cm | 3 | 3 | 3 | 3 |
| Bioirrigation | $\alpha_0$ | $yr^{-1}$ | 0 | 0 | 0 | 0 |

| | | | | | | |
|---|---|---|---|---|---|---|
| coeffcient | | | | | | |
| $R_1$ $O_2$ inhibition concentration | $h_1$ | μM | 10 | 10 | 10 | 10 |
| $R_2$ $NO_3^-$ inhibition concentration | $h_2$ | μM | 1 | 10 | 1 | 1 |
| Denitrification deceleration constant | $k_6$ | -- | 0.2 | 0.3 | 0.8 | 0.8 |
| MnO2 reduction deceleration constant | $k_7$ | -- | 0.018 | 0.018 | 0.08 | 0.035 |
| Porosity at sediment surface | $\varphi_0$ | -- | 0.8 | 0.8 | 0.65 | 0.8 |
| Porosity at infinite depth | $\varphi_\infty$ | -- | 0.55 | 0.6 | 0.55 | 0.6 |
| Porosity attenuation coefficient | $\alpha_0$ | $cm^{-1}$ | 0.01 | 0.01 | 0.01 | 0.01 |

**Table S4. Summary statistics of Candidatus *Scalindua* genomes**

| | ***Ca.* S. sediminis (this study)** | ***Ca.* S. rubra** | ***Ca.* S. profunda** | ***Ca.* S. brodae** | ***Ca.* S. japonica** | ***Ca.* S. AMX11** |
|---|---|---|---|---|---|---|
| Completeness* | 95.5% | 92.5% | 94.2% | 92.7% | 95.5% | 96.6% |
| Contamination* | 4.6% | 5.1% | 5.4% | 2.3% | 3.4% | 4.6% |
| Strain heterogeneity* | 0% | 0% | 75% | 0% | 0% | 25% |
| Total length (base pairs) | 2,955,644 | 5,194,263 | 4,176,727 | 4,084,168 | 4,812,853 | 4,593,657 |
| GC content | 38.3% | 37.3% | 40.3% | 39.6% | 38.8% | 41.1% |
| Number of scaffolds | 71 | 443 | 3053 | 282 | 47 | 121 |
| Number of contigs | 83 | 443 | 4756 | 282 | 47 | 121 |
| N50 of contigs | 70,308 | 22,837 | 1,173 | 33,252 | 219,109 | 92,628 |
| Number of coding sequences† | 2,879 | 5,482 | 4,714 | 4,178 | 4,343 | 4095 |
| Coding density | 84.1% | 80.1% | 99.4% | 83.9% | 84.1% | 81.6% |
| Average coverage | 116.7 | -- | -- | -- | -- | -- |
| iRep | 1.32 | -- | -- | -- | -- | -- |

*Based on lineage-specific marker sets determined with CheckM. †Inferred with Prodigal (ref.). ‡Estimated from the proportion of reads mapped to the genome. --, No data.

**Table S5. Occurrence of *Ca.* Scalindua sediminis-like bacteria in natural environments**

| Sample ID in NCBI | # Total sequences | Sample Origin | 99% similarity | 97% similarity | % matching reads |
|---|---|---|---|---|---|
| SRR6419741 | 85617 | AMOR subseafloor sediment | 0 | 3592 | 4.20 |
| SRR6419772 | 121701 | AMOR subseafloor sediment | 0 | 4544 | 3.73 |
| SRR6419774 | 67467 | AMOR subseafloor sediment | 0 | 2143 | 3.18 |
| SRR6419746 | 72258 | AMOR subseafloor sediment | 0 | 1925 | 2.66 |
| SRR6419794 | 114471 | AMOR subseafloor sediment | 0 | 2819 | 2.46 |
| SRR6419775 | 99355 | AMOR subseafloor sediment | 0 | 2442 | 2.46 |
| ERR1360111 | 30458 | South China Sea sediments | 0 | 724 | 2.38 |
| SRR2048121 | 13234 | Equatorial Pacific sediment | 0 | 305 | 2.30 |
| SRR6419787 | 89402 | AMOR subseafloor sediment | 0 | 1946 | 2.18 |
| SRR6419747 | 108128 | AMOR subseafloor sediment | 0 | 2317 | 2.14 |
| ERR1360114 | 13433 | South China Sea sediments | 0 | 284 | 2.11 |

| | | | | | |
|---|---|---|---|---|---|
| SRR6419785 | 154199 | AMOR subseafloor sediment | 0 | 3225 | 2.09 |
| SRR1824401 | 54338 | marine methane seeps sediment | 0 | 1085 | 2.00 |
| SRR6419784 | 79020 | AMOR subseafloor sediment | 0 | 1389 | 1.76 |
| SRR6419788 | 126025 | AMOR subseafloor sediment | 0 | 2187 | 1.74 |
| SRR6419786 | 118833 | AMOR subseafloor sediment | 0 | 2055 | 1.73 |
| SRR1824369 | 55720 | marine methane seeps sediment | 0 | 911 | 1.63 |
| SRR6419740 | 139454 | AMOR subseafloor sediment | 0 | 2186 | 1.57 |
| SRR396763 | 3911 | AMOR sediments | 57 | 58 | 1.48 |
| SRR6419789 | 124052 | AMOR subseafloor sediment | 0 | 1813 | 1.46 |
| ERR1360103 | 53491 | South China Sea sediments | 699 | 762 | 1.42 |
| SRR1824399 | 33640 | marine methane seeps sediment | 0 | 465 | 1.38 |
| SRR2071711 | 9964 | Saanich Inlet | 0 | 111 | 1.11 |
| SRR1824386 | 60162 | marine methane seeps sediment | 603 | 606 | 1.01 |
| SRR1824373 | 48295 | marine methane seeps sediment | 0 | 421 | 0.87 |
| SRR6419743 | 121923 | AMOR subseafloor sediment | 0 | 1052 | 0.86 |
| SRR6419773 | 123348 | AMOR subseafloor sediment | 0 | 941 | 0.76 |
| SRR1824392 | 44305 | marine methane seeps sediment | 0 | 319 | 0.72 |
| SRR1736202 | 4754 | shallow-water pockmarks sediments | 0 | 34 | 0.72 |
| SRR1824398 | 66227 | marine methane seeps sediment | 0 | 449 | 0.68 |
| SRR6419745 | 139235 | AMOR subseafloor sediment | 0 | 873 | 0.63 |
| SRR1824377 | 29292 | marine methane seeps sediment | 0 | 177 | 0.60 |
| SRR6419769 | 87195 | AMOR subseafloor sediment | 0 | 522 | 0.60 |
| SRR6419783 | 102145 | AMOR subseafloor sediment | 584 | 607 | 0.59 |
| SRR1736199 | 5575 | shallow-water pockmarks sediments | 0 | 33 | 0.59 |
| SRR1824376 | 48908 | marine methane seeps sediment | 0 | 277 | 0.57 |
| SRR6419742 | 104151 | AMOR subseafloor sediment | 0 | 554 | 0.53 |
| SRR1824394 | 53071 | marine methane seeps sediment | 0 | 280 | 0.53 |
| SRR978583 | 16173 | submarine sediments 1 mbsf | 0 | 85 | 0.53 |
| SRR2048149 | 11777 | Equatorial Pacific sediment | 0 | 59 | 0.50 |
| SRR1824395 | 75205 | marine methane seeps sediment | 0 | 361 | 0.48 |
| SRR1824357 | 57359 | marine methane seeps sediment | 0 | 272 | 0.47 |
| SRR1736201 | 4409 | shallow-water pockmarks sediments | 0 | 19 | 0.43 |
| ERR1360104 | 45322 | South China Sea sediments | 0 | 192 | 0.42 |
| SRR978642 | 9177 | submarine sediments 1 mbsf | 0 | 38 | 0.41 |
| SRR1736198 | 3960 | shallow-water pockmarks sediments | 0 | 16 | 0.40 |
| SRR6419753 | 111907 | AMOR subseafloor sediment | 0 | 448 | 0.40 |
| SRR1295391 | 33116 | marine sediment close to Hakon Mosby Mud Volcano | 130 | 132 | 0.40 |
| SRR1824400 | 49815 | marine methane seeps sediment | 0 | 192 | 0.39 |
| SRR978587 | 8381 | submarine sediments 1 mbsf | 0 | 31 | 0.37 |
| SRR1824360 | 67479 | marine methane seeps sediment | 0 | 248 | 0.37 |
| SRR6419768 | 88607 | AMOR subseafloor sediment | 0 | 325 | 0.37 |
| SRR1824350 | 56190 | marine methane seeps sediment | 0 | 205 | 0.36 |
| SRR1824383 | 48085 | marine methane seeps sediment | 0 | 159 | 0.33 |
| SRR978588 | 9129 | submarine sediments 1 mbsf | 0 | 30 | 0.33 |

| | | | | | |
|---|---|---|---|---|---|
| SRR6419782 | 116403 | AMOR subseafloor sediment | 0 | 364 | 0.31 |
| SRR1824406 | 49359 | marine methane seeps sediment | 0 | 151 | 0.31 |
| SRR1824387 | 51023 | marine methane seeps sediment | 147 | 156 | 0.31 |
| SRR1824388 | 77220 | marine methane seeps sediment | 0 | 212 | 0.27 |
| SRR978585 | 12796 | submarine sediments 1 mbsf | 0 | 33 | 0.26 |
| SRR1824402 | 45019 | marine methane seeps sediment | 0 | 116 | 0.26 |
| SRR1824381 | 39793 | marine methane seeps sediment | 0 | 101 | 0.25 |
| ERR1360113 | 18750 | South China Sea sediments | 46 | 46 | 0.25 |
| SRR978584 | 15164 | submarine sediments 1 mbsf | 0 | 37 | 0.24 |
| SRR978582 | 15549 | submarine sediments 1 mbsf | 0 | 37 | 0.24 |
| SRR1824384 | 45344 | marine methane seeps sediment | 0 | 105 | 0.23 |
| ERR951487 | 4914 | cold-water coral mounds sediments | 11 | 11 | 0.22 |
| SRR6419755 | 142593 | AMOR subseafloor sediment | 0 | 317 | 0.22 |
| SRR1824411 | 45798 | marine methane seeps sediment | 0 | 100 | 0.22 |
| SRR1824355 | 32690 | marine methane seeps sediment | 0 | 69 | 0.21 |
| SRR1736200 | 4773 | shallow-water pockmarks sediments | 0 | 10 | 0.21 |
| SRR1559242 | 33425 | East China Sea sediments | 0 | 70 | 0.21 |
| SRR978684 | 9565 | submarine sediments | 0 | 20 | 0.21 |
| SRR1559234 | 101902 | East China Sea sediments | 0 | 213 | 0.21 |
| SRR978586 | 7789 | submarine sediments | 0 | 16 | 0.21 |
| SRR1824385 | 65897 | marine methane seeps sediment | 0 | 133 | 0.20 |
| SRR1824375 | 52991 | marine methane seeps sediment | 0 | 100 | 0.19 |
| SRR1824353 | 49294 | marine methane seeps sediment | 0 | 91 | 0.18 |
| SRR1824391 | 51510 | marine methane seeps sediment | 0 | 94 | 0.18 |
| SRR6419752 | 123591 | AMOR subseafloor sediment | 0 | 222 | 0.18 |
| SRR6419757 | 122074 | AMOR subseafloor sediment | 0 | 217 | 0.18 |
| SRR1824389 | 43990 | marine methane seeps sediment | 0 | 78 | 0.18 |
| SRR1824382 | 59423 | marine methane seeps sediment | 0 | 104 | 0.18 |
| SRR1559238 | 27534 | East China Sea sediments | 0 | 48 | 0.17 |
| SRR1824379 | 60871 | marine methane seeps sediment | 0 | 105 | 0.17 |
| SRR1824351 | 37082 | marine methane seeps sediment | 0 | 62 | 0.17 |
| SRR6419771 | 106536 | AMOR subseafloor sediment | 0 | 177 | 0.17 |
| SRR1824397 | 48011 | marine methane seeps sediment | 77 | 77 | 0.16 |
| SRR6396235 | 832341 | Atlantic Surface Sediment | 0 | 1314 | 0.16 |
| SRR6419781 | 133185 | AMOR subseafloor sediment | 201 | 209 | 0.16 |
| SRR1647975 | 46294 | marine methane seeps sediment | 0 | 71 | 0.15 |
| SRR1824352 | 58719 | marine methane seeps sediment | 0 | 82 | 0.14 |
| SRR1797781 | 8683 | AMOR sediment close to LC | 9 | 12 | 0.14 |
| SRR1824359 | 46843 | marine methane seeps sediment | 0 | 64 | 0.14 |
| ERR1360106 | 15634 | South China Sea sediments | 0 | 21 | 0.13 |
| SRR1824396 | 55946 | marine methane seeps sediment | 0 | 74 | 0.13 |
| SRR6419756 | 109310 | AMOR subseafloor sediment | 0 | 144 | 0.13 |
| SRR1824390 | 35707 | marine methane seeps sediment | 0 | 47 | 0.13 |
| SRR1559243 | 47047 | East China Sea sediments | 0 | 56 | 0.12 |

| | | | | | |
|---|---|---|---|---|---|
| SRR6419754 | 125536 | AMOR subseafloor sediment | 0 | 136 | 0.11 |
| SRR6419770 | 81397 | AMOR subseafloor sediment | 0 | 76 | 0.09 |
| SRR1559241 | 32524 | East China Sea sediments | 0 | 29 | 0.09 |
| SRR1559240 | 22172 | East China Sea sediments | 0 | 18 | 0.08 |
| ERR1360115 | 31048 | South China Sea sediments | 0 | 25 | 0.08 |
| SRR1824339 | 53172 | marine methane seeps sediment | 0 | 39 | 0.07 |
| SRR1559236 | 39181 | East China Sea sediments | 0 | 28 | 0.07 |
| ERR1685370 | 77502 | Faroe Shetland Channel Deep-sea sediment | 0 | 54 | 0.07 |
| SRR1824405 | 54744 | marine methane seeps sediment | 0 | 38 | 0.07 |
| SRR1559259 | 36427 | East China Sea sediments | 0 | 25 | 0.07 |
| SRR1559244 | 25223 | East China Sea sediments | 0 | 16 | 0.06 |
| DRR060154 | 24327 | East China Sea coastal sediments | 0 | 15 | 0.06 |
| SRR1824378 | 72617 | marine methane seeps sediment | 0 | 38 | 0.05 |
| SRR1824347 | 42890 | marine methane seeps sediment | 0 | 22 | 0.05 |
| ERR2016813 | 121609 | Barents Sea sediment 1400 m | 60 | 60 | 0.05 |
| SRR1119244 | 35383 | GOM oil plume | 0 | 16 | 0.05 |

**Table S6. Annotation of *Ca.* S. sediminis genes discussed in this study.**

| Pathway | gene_callers_id | contig | start | stop | Gene/Product | Function |
|---|---|---|---|---|---|---|
| Complex I | 1490 | c_000000000013 | 68802 | 69159 | nuoA | NADH-quinone oxidoreductase subunit A |
| Complex I | 1491 | c_000000000013 | 69149 | 69635 | nuoB | NADH-quinone oxidoreductase subunit B |
| Complex I | 2505 | c_000000000040 | 13157 | 14609 | nuoN | NADH-quinone oxidoreductase subunit N |
| Complex I | 2506 | c_000000000040 | 14671 | 16243 | nuoM | NADH-quinone oxidoreductase subunit M |
| Complex I | 2507 | c_000000000040 | 16524 | 18444 | nuoL | NADH-quinone oxidoreductase subunit L |
| Complex I | 2508 | c_000000000040 | 18481 | 18784 | nuoK | NADH-quinone oxidoreductase subunit 11 |
| Complex I | 2509 | c_000000000040 | 18788 | 19313 | nuoJ | NADH-quinone oxidoreductase subunit J |
| Complex I | 2510 | c_000000000040 | 19334 | 19754 | nuoI | NADH-quinone oxidoreductase subunit 9 |
| Complex I | 747 | c_000000000005 | 29512 | 30208 | nqrC | Na(+)-translocating NADH-quinone reductase subunit C |
| Complex I | 748 | c_000000000005 | 30204 | 31623 | nqrE | Na(+)-translocating NADH-quinone reductase subunit E |
| Complex I | 749 | c_000000000005 | 31664 | 33299 | nqrF | Na(+)-translocating NADH-quinone reductase subunit F |
| Complex II | 583 | c_000000000004 | 7010 | 7358 | sdhD | Succinate dehydrogenase hydrophobic membrane anchor protein |
| Complex II | 584 | c_000000000004 | 7377 | 7770 | sdhC | Succinate dehydrogenase 2 membrane subunit SdhC |
| Complex II | 585 | c_000000000004 | 7890 | 9681 | sdhA | Fumarate reductase flavoprotein subunit |
| Complex II | 586 | c_000000000004 | 9704 | 10409 | sdhB | Succinate dehydrogenase iron-sulfur subunit |
| Complex III | 158 | c_000000000001 | 168102 | 168948 | ndhI_1 | NAD(P)H-quinone oxidoreductase subunit I chloroplastic |
| Complex III | 160 | c_000000000001 | 169788 | 171834 | ndhI_2 | NAD(P)H-quinone oxidoreductase subunit I chloroplastic |
| Complex III | 161 | c_000000000001 | 172088 | 172958 | sdhE | 8-methylmenaquinol:fumarate reductase membrane anchor subuni[illegible] |
| Complex III | 162 | c_000000000001 | 172966 | 173521 | sdhB_1 | 8-methylmenaquinol:fumarate reductase iron-sulfur subunit |
| Complex III | 533 | c_000000000003 | 118126 | 118837 | petB_1 | Cytochrome b6 |
| Complex III | 534 | c_000000000003 | 118859 | 119258 | petD | Cytochrome b6-f complex subunit 4 |
| Complex III | 965 | c_000000000007 | 24814 | 25585 | petB_2 | Cytochrome b6 |
| Complex III | 1359 | c_000000000012 | 12782 | 13373 | cycA1 | Cytochrome c-554 |
| Complex III | 1476 | c_000000000013 | 53972 | 55307 | psbV | Cytochrome c-550 |

| | | | | | | |
|---|---|---|---|---|---|---|
| Complex III | 1843 | c_000000000020 | 44893 | 45745 | petB_3 | Cytochrome b6 |
| Complex III | 1844 | c_000000000020 | 45898 | 46432 | petC_2 | Cytochrome b6-f complex iron-sulfur subunit |
| Complex III | 2096 | c_000000000026 | 8080 | 8854 | petB_4 | Cytochrome b6 |
| Complex III | 2098 | c_000000000026 | 10777 | 11296 | petC_3 | Cytochrome b6-f complex iron-sulfur subunit |
| Complex IV | 1146 | c_000000000009 | 18405 | 18747 | fixP_1 | Cbb3-type cytochrome c oxidase subunit FixP |
| Complex IV | 1147 | c_000000000009 | 18866 | 19214 | fixP_2 | Cbb3-type cytochrome c oxidase subunit FixP |
| Complex V | 940 | c_000000000007 | 1142 | 1694 | atpH | F1 sector of membrane-bound ATP synthase, delta subunit |
| Complex V | 941 | c_000000000007 | 1727 | 3251 | atpA | F1 sector of membrane-bound ATP synthase, alpha subunit |
| Complex V | 942 | c_000000000007 | 3292 | 4189 | atpG | F1 sector of membrane-bound ATP synthase, gamma subunit |
| Complex V | 943 | c_000000000007 | 4305 | 5718 | atpD | F1 sector of membrane-bound ATP synthase, beta subunit |
| Complex V | 944 | c_000000000007 | 5782 | 6049 | atpC | F1 sector of membrane-bound ATP synthase, epsilon subunit |
| Complex V | 2631 | c_000000000047 | 827 | 1397 | atpB | F0 sector of membrane-bound ATP synthase, subunit a |
| Complex V | 2632 | c_000000000047 | 1444 | 1684 | atpE | F0 sector of membrane-bound ATP synthase, subunit c |
| Complex V | 2633 | c_000000000047 | 1772 | 2276 | atpF | F0 sector of membrane-bound ATP synthase, subunit b |
| TCA cycle | 582 | c_000000000004 | 6030 | 6951 | mdh | Malate dehydrogenase |
| TCA cycle | 583 | c_000000000004 | 7010 | 7358 | sdhD | Succinate dehydrogenase hydrophobic membrane anchor protein |
| TCA cycle | 584 | c_000000000004 | 7377 | 7770 | sdhC | Succinate dehydrogenase 2 membrane subunit SdhC |
| TCA cycle | 585 | c_000000000004 | 7890 | 9681 | sdhA | Fumarate reductase flavoprotein subunit |
| TCA cycle | 586 | c_000000000004 | 9704 | 10409 | sdhB | Succinate dehydrogenase iron-sulfur subunit |
| TCA cycle | 628 | c_000000000004 | 47716 | 48847 | citZ | Citrate synthase |
| TCA cycle | 844 | c_000000000006 | 7612 | 8359 | korB_1 | 2-oxoglutarate oxidoreductase subunit KorB |
| TCA cycle | 845 | c_000000000006 | 8381 | 9473 | korA_1 | 2-oxoglutarate oxidoreductase subunit KorA |
| TCA cycle | 1101 | c_000000000008 | 71801 | 72539 | korB_2 | 2-oxoglutarate oxidoreductase subunit KorB |
| TCA cycle | 1102 | c_000000000008 | 72535 | 73555 | korA_2 | 2-oxoglutarate oxidoreductase subunit KorA |
| TCA cycle | 1104 | c_000000000008 | 73855 | 74722 | sucD | Succinate--CoA ligase [ADP-forming] subunit alpha |

| | | | | | | |
|---|---|---|---|---|---|---|
| TCA cycle | 1105 | c_000000000008 | 74725 | 75889 | sucC | Succinate--CoA ligase [ADP-forming] subunit beta |
| TCA cycle | 1812 | c_000000000020 | 16830 | 18075 | icd | Isocitrate dehydrogenase [NADP] |
| TCA cycle | 1142 | c_000000000009 | 15196 | 16627 | fumC | Fumarate hydratase (fumarase C), aerobic Class II |
| Hydrogen metabolism | | | | | | |
| Hydrogen metabolism | 1555 | c_000000000015 | 2364 | 3603 | hupR1_1 | Hydrogenase transcriptional regulatory protein hupR1 |
| Hydrogen metabolism | 2025 | c_000000000024 | 24028 | 25345 | hupR1_2 | Hydrogenase transcriptional regulatory protein hupR1 |
| Hydrogen metabolism | 2726 | c_000000000054 | 2734 | 4036 | hupR1_3 | Hydrogenase transcriptional regulatory protein hupR1 |
| Hydrogen metabolism | 2524 | c_000000000041 | 12450 | 12936 | hndA | NADP-reducing hydrogenase subunit HndA |
| Hydrogen metabolism | 2525 | c_000000000041 | 12939 | 14577 | hndC | NADP-reducing hydrogenase subunit HndC |
| **Miscellaneous** | | | | | | |
| **Amino acid transport** | | | | | | |
| Branched AA Transport | 1166 | c_000000000009 | 39168 | 40839 | livH | High-affinity branched-chain amino acid transport system permeas |
| Branched AA Transport | 1169 | c_000000000009 | 43036 | 43786 | livF | High-affinity branched-chain amino acid transport ATP-binding pr |
| Peptide Transport | 1710 | c_000000000017 | 43102 | 44782 | cstA | Peptide transporter CstA |
| Phospholipid Transport | 791 | c_000000000005 | 75513 | 76314 | mlaE | putative phospholipid ABC transporter permease protein MlaE |
| Phospholipid Transport | 792 | c_000000000005 | 76313 | 77054 | mkl | putative ribonucleotide transport ATP-binding protein mkl |
| Phospholipid Transport | 793 | c_000000000005 | 77201 | 78155 | mlaD | putative phospholipid ABC transporter-binding protein MlaD |
| Arsenic Resistance | 1365 | c_000000000012 | 17656 | 18793 | acr3 | Arsenical-resistance protein Acr3 |
| Arsenic Resistance | 1366 | c_000000000012 | 18840 | 19209 | arsC_2 | Arsenate reductase |
| Arsenic Resistance | 932 | c_000000000006 | 108529 | 108970 | arsC_1 | Arsenate reductase |
| Carbonic Anhydrases | 2531 | c_000000000042 | 2136 | 2733 | can | Carbonic anhydrase 2 |
| Co/Zn/Cd Export | 1492 | c_000000000014 | 383 | 1247 | cbiM | Cobalt transport protein CbiM |
| Co/Zn/Cd Export | 1493 | c_000000000014 | 1353 | 1776 | CbiN | Cobalt transport protein CbiN |
| Co/Zn/Cd Export | 1494 | c_000000000014 | 1775 | 2561 | cbiQ | Cobalt transport protein CbiQ |

| | | | | | | |
|---|---|---|---|---|---|---|
| Co/Zn/Cd Export | 1062 | c_000000000008 | 26528 | 27614 | czcB_1 | Cobalt-zinc-cadmium resistance protein CzcB |
| Co/Zn/Cd Export | 1673 | c_000000000017 | 5444 | 6641 | czcB_2 | Cobalt-zinc-cadmium resistance protein CzcB |
| Co/Zn/Cd Export | 1674 | c_000000000017 | 6637 | 9766 | czcA_2 | Cobalt-zinc-cadmium resistance protein CzcA |
| Co/Zn/Cd Export | 1675 | c_000000000017 | 9762 | 11052 | czcC_2 | Cobalt-zinc-cadmium resistance protein CzcC |
| Co/Zn/Cd Export | 1908 | c_000000000022 | 13676 | 16805 | czcA_3 | Cobalt-zinc-cadmium resistance protein CzcA |
| Co/Zn/Cd Export | 226 | c_000000000001 | 243555 | 246666 | czcA_1 | Cobalt-zinc-cadmium resistance protein CzcA |
| Co/Zn/Cd Export | 537 | c_000000000003 | 121943 | 122555 | czcD_2 | Cadmium cobalt and zinc/H(+)-K(+) antiporter |
| Co/Zn/Cd Export | 214 | c_000000000001 | 229660 | 230611 | czcD_1 | Cadmium cobalt and zinc/H(+)-K(+) antiporter |
| CRISPR/CAS | 1303 | c_000000000011 | 27651 | 27939 | cas2 | CRISPR-associated endoribonuclease Cas2 |
| CRISPR/CAS | 1304 | c_000000000011 | 27948 | 28980 | cas1 | CRISPR-associated endonuclease Cas1 |
| CRISPR/CAS | 1305 | c_000000000011 | 29054 | 29693 | cas4 | CRISPR-associated endonuclease Cas4 |
| CRISPR/CAS | 1307 | c_000000000011 | 30264 | 31170 | cas7/cas2 | CRISPR-associated endonuclease Cas7/Cas2 |
| CRISPR/CAS | 1308 | c_000000000011 | 31199 | 33365 | cas8c/csd1 | CRISPR-associated endonuclease Cas8c/Csd1 |
| CRISPR/CAS | 1309 | c_000000000011 | 33361 | 34087 | cas5d | CRISPR pre-crRNA endoribonuclease Cas5d |
| Cytochrome c biogenesis | 1435 | c_000000000013 | 13885 | 14779 | ccsA_1 | Cytochrome c biogenesis protein CcsA |
| Cytochrome c biogenesis | 2339 | c_000000000033 | 4712 | 5597 | ccsA_2 | Cytochrome c biogenesis protein CcsA |
| Cytochrome c biogenesis | 2340 | c_000000000033 | 5690 | 7568 | ccs1 | Cytochrome c biogenesis protein Ccs1 |
| Cytochrome c biogenesis | 2706 | c_000000000052 | 138 | 900 | ccsA_3 | Cytochrome c biogenesis protein CcsA |
| Magnesium Transport | 246 | c_000000000001 | 271842 | 273192 | mgtE | Magnesium transporter MgtE |
| Sodium/Proton Antiporter | 2639 | c_000000000047 | 5931 | 6414 | mnhE1 | Na(+)/H(+) antiporter subunit E1 |
| Sodium/Proton Antiporter | 2640 | c_000000000047 | 6406 | 6661 | mrpF | Na(+)/H(+) antiporter subunit F |
| Sodium/Proton Antiporter | 2642 | c_000000000047 | 7259 | 7571 | mrpG | Na(+)/H(+) antiporter subunit G |
| Sodium/Proton Antiporter | 2643 | c_000000000047 | 7633 | 8212 | mrpA_1 | Na(+)/H(+) antiporter subunit A |
| Sodium/Proton Antiporter | 2644 | c_000000000047 | 8435 | 8873 | mrpB | Na(+)/H(+) antiporter subunit B |

| | | | | | | |
|---|---|---|---|---|---|---|
| Sodium/Proton Antiporter | 2645 | c_000000000047 | 8879 | 9305 | mrpC | Na(+)/H(+) antiporter subunit C |
| Sodium/Proton Antiporter | 2646 | c_000000000047 | 9306 | 10833 | mrpD | Na(+)/H(+) antiporter subunit D |
| Sodium/Proton Antiporter | 2650 | c_000000000047 | 13296 | 15108 | mrpA_2 | Na(+)/H(+) antiporter subunit A |
| Sodium/Proton Antiporter | 741 | c_000000000005 | 21464 | 22721 | gerN_1 | Na(+)/H(+)-K(+) antiporter GerN |
| Sodium/Proton Antiporter | 742 | c_000000000005 | 22738 | 24280 | gerN_2 | Na(+)/H(+)-K(+) antiporter GerN |
| Sodium Extrusion | 1351 | c_000000000012 | 4390 | 4921 | ntpK | V-type sodium ATPase subunit K |
| Sodium Extrusion | 1352 | c_000000000012 | 5127 | 5694 | atpE_1 | V-type proton ATPase subunit E |
| Sodium Extrusion | 1353 | c_000000000012 | 5892 | 6993 | ntpC | V-type sodium ATPase subunit C |
| Sodium Extrusion | 1354 | c_000000000012 | 6985 | 7300 | ntpG | V-type sodium ATPase subunit G |
| Sodium Extrusion | 1355 | c_000000000012 | 7774 | 9541 | ntpA | V-type sodium ATPase catalytic subunit A |
| Sodium Extrusion | 1356 | c_000000000012 | 9558 | 10938 | ntpB | V-type sodium ATPase subunit B |
| Sodium Extrusion | 1357 | c_000000000012 | 10934 | 11591 | ntpD | V-type sodium ATPase subunit D |
| ATP Synthesis | 2631 | c_000000000047 | 827 | 1397 | atpB | ATP synthase subunit a |
| ATP Synthesis | 2632 | c_000000000047 | 1444 | 1684 | atpE | ATP synthase subunit c sodium ion specific |
| ATP Synthesis | 2633 | c_000000000047 | 1772 | 2276 | atpF | ATP synthase subunit b |
| Phosphate Transport | 1370 | c_000000000012 | 22476 | 23394 | pstS | Phosphate-binding protein PstS |
| Phosphate Transport | 1371 | c_000000000012 | 23390 | 24275 | pstC1 | Phosphate transport system permease protein PstC 1 |
| Phosphate Transport | 1372 | c_000000000012 | 24271 | 25126 | pstA1 | Phosphate transport system permease protein PstA 1 |
| Phosphate Transport | 1373 | c_000000000012 | 25146 | 25899 | pstB3 | Phosphate import ATP-binding protein PstB 3 |
| Phosphate Transport | 1374 | c_000000000012 | 25963 | 26635 | phoU | Phosphate-specific transport system accessory protein PhoU |
| Phosphate Transport | 1367 | c_000000000012 | 19201 | 19882 | phoB_2 | Phosphate regulon transcriptional regulatory protein PhoB |
| Phosphate Transport | 1660 | c_000000000016 | 51870 | 52998 | pitA_1 | Low-affinity inorganic phosphate transporter 1 |
| Phosphate Transport | 1686 | c_000000000017 | 21307 | 22732 | pitA_2 | Low-affinity inorganic phosphate transporter 1 |
| Formate Transport | 956 | c_000000000007 | 16707 | 17703 | focA_1 | putative formate transporter 1 |

| | | | | | | |
|---|---|---|---|---|---|---|
| Formate Transport | 957 | c_000000000007 | 17715 | 18618 | focA_2 | putative formate transporter 1 |
| RND family/Heavy metal/cation/multidrg Efflux | 553 | c_000000000003 | 134790 | 138762 | cusA | Cation efflux system protein CusA |
| RND family/Heavy metal/cation/multidrg Efflux | 555 | c_000000000003 | 139054 | 141211 | cusB | Cation efflux system protein CusB |
| RND family/Heavy metal/cation/multidrg Efflux | 2378 | c_000000000034 | 16357 | 17251 | fieF_1 | Ferrous-iron efflux pump FieF |
| RND family/Heavy metal/cation/multidrg Efflux | 2379 | c_000000000034 | 17280 | 18504 | | putative transporter |
| RND family/Heavy metal/cation/multidrg Efflux | 2386 | c_000000000034 | 23052 | 24003 | fieF_2 | Ferrous-iron efflux pump FieF |
| RND family/Heavy metal/cation/multidrg Efflux | 615 | c_000000000004 | 35616 | 37245 | bepC | Outer membrane efflux protein BepC |
| RND family/Heavy metal/cation/multidrg Efflux | 225 | c_000000000001 | 242316 | 243417 | ttgG | Toluene efflux pump periplasmic linker protein TtgG |
| RND family/Heavy metal/cation/multidrg Efflux | 1061 | c_000000000008 | 22756 | 25954 | mdtC | Multidrug resistance protein MdtC |
| RND family/Heavy metal/cation/multidrg Efflux | 1094 | c_000000000008 | 61999 | 63760 | yheI | putative multidrug resistance ABC transporter ATP-binding/perme |
| RND family/Heavy metal/cation/multidrg Efflux | 1901 | c_000000000022 | 5994 | 9237 | mdtB | Multidrug resistance protein MdtB |
| RND family/Heavy | 132 | c_000000000001 | 135473 | 137339 | | Putative multidrug export ATP-binding/permease protein |

| | | | | | | |
|---|---|---|---|---|---|---|
| metal/cation/multidrg Efflux | | | | | | |
| Ferredoxin | 542 | c_000000000003 | 126765 | 126954 | | Ferredoxin |
| Ferredoxin | 2487 | c_000000000039 | 16592 | 16781 | | Ferredoxin |
| Ferredoxin | 729 | c_000000000005 | 9733 | 10009 | fixX | Ferredoxin-like protein FixX |
| Ferredoxin | 1188 | c_000000000009 | 62744 | 63842 | fdxA | 4Fe-4S ferredoxin FdxA |
| Ferredoxin | 1243 | c_000000000010 | 30383 | 32870 | napG_1 | Ferredoxin-type protein NapG |
| Ferredoxin | 2574 | c_000000000044 | 12634 | 13267 | napG_2 | Ferredoxin-type protein NapG |
| Sec Pathway | 88 | c_000000000001 | 91600 | 92983 | secY | Protein translocase subunit SecY |
| Sec Pathway | 119 | c_000000000001 | 117259 | 117718 | secE | Protein translocase subunit SecE |
| Sec Pathway | 1426 | c_000000000013 | 5053 | 5374 | yajC | Sec translocon accessory complex subunit YajC |
| Sec Pathway | 1427 | c_000000000013 | 5414 | 8762 | secD | Protein translocase subunit SecD |
| Sec Pathway | 1610 | c_000000000015 | 61368 | 63984 | secA | Protein translocase subunit SecA |
| TAT secretion | 436 | c_000000000003 | 14415 | 14670 | tatA_1 | Sec-independent protein translocase protein TatA |
| TAT secretion | 904 | c_000000000006 | 76488 | 76695 | tatA_2 | Sec-independent protein translocase protein TatA |
| TAT secretion | 1255 | c_000000000010 | 44947 | 46276 | tatC2 | Sec-independent protein translocase protein TatCy |
| Type II secretion | 8 | c_000000000001 | 9714 | 10386 | pulG_1 | Type II secretion system protein G |
| Type II secretion | 910 | c_000000000006 | 79444 | 80695 | epsF | Type II secretion system protein F |
| Type II secretion | 911 | c_000000000006 | 80931 | 82662 | epsE_2 | Type II secretion system protein E |
| Type II secretion | 912 | c_000000000006 | 82674 | 84378 | epsE_3 | Type II secretion system protein E |
| Type II secretion | 1716 | c_000000000017 | 52256 | 52574 | epsG | Type II secretion system protein G |
| Type II secretion | 1717 | c_000000000017 | 52731 | 54399 | xpsE | Type II secretion system protein E |
| Type II secretion | 1762 | c_000000000018 | 46001 | 46676 | pulG_2 | Type II secretion system protein G |
| **Nitrogen metabolism** | | | | | | |

| | | | | | | |
|---|---|---|---|---|---|---|
| Ammonium transport | 1206 | c_000000000009 | 85170 | 86505 | nrgA | Ammonium transporter |
| Ammonium transport | 1201 | c_000000000009 | 79299 | 80850 | amtB_2 | Ammonia channel |
| Ammonium transport | 1203 | c_000000000009 | 81642 | 83028 | amtB | Ammonia channel |
| Nitrite oxidation | 2567 | c_000000000044 | 1859 | 2810 | nxrC | nitrite oxidoreductase gamma subunit |
| Nitrite oxidation | 2568 | c_000000000044 | 2857 | 4120 | nxrB | nitrite oxidoreductase beta subunit |
| Nitrite oxidation | 2570 | c_000000000044 | 5032 | 5806 | nxrD | nitrite oxidoreductase Chaperone protein delta subunit |
| Nitrite oxidation | 2572 | c_000000000044 | 6828 | 10317 | nxrA | nitrite oxidoreductase alpha subunit |
| Urea metabolism | 1172 | c_000000000009 | 46910 | 47210 | ureA | Urease subunit gamma |
| Urea metabolism | 1173 | c_000000000009 | 47302 | 47674 | ureB | Urease subunit beta |
| Urea metabolism | 1174 | c_000000000009 | 47711 | 49427 | ureC | Urease subunit alpha |
| Urea metabolism | 1176 | c_000000000009 | 50212 | 50767 | ureE | Urease accessory protein UreE |
| Urea metabolism | 1177 | c_000000000009 | 50715 | 51525 | ureF | Urease accessory protein UreF |
| Urea metabolism | 1178 | c_000000000009 | 51645 | 52269 | ureG | Urease accessory protein UreG |
| Urea metabolism | 1179 | c_000000000009 | 52281 | 53220 | ureD1 | Urease accessory protein UreD |
| Urea metabolism | 1180 | c_000000000009 | 53261 | 54830 | groL_2 | 60 kDa chaperonin |
| Cyanate metabolism | 2085 | c_000000000026 | 138 | 591 | cynS_1 | Cyanate hydratase |
| Cyanate metabolism | 2773 | c_000000000061 | 3077 | 3530 | cynS_2 | Cyanate hydratase |
| Hydrazine metabolism | 1033 | c_000000000007 | 95308 | 96445 | hzsB | Hydrazine synthase subunit beta |
| Hydrazine metabolism | 1034 | c_000000000007 | 97201 | 98230 | hzsA | Hydrazine synthase subunit gamma |
| Hydrazine metabolism | 1671 | c_000000000017 | 1404 | 2607 | hzsB_2 | Hydrazine synthase subunit beta |
| Hydrazine metabolism | 1789 | c_000000000019 | 29192 | 31211 | hzsB_3 | Hydrazine synthase subunit beta |
| Hydrazine metabolism | 2149 | c_000000000027 | 26133 | 27270 | hzsB | Hydrazine synthase subunit beta |
| Hydrazine metabolism | 2150 | c_000000000027 | 27332 | 28358 | hzsC | Hydrazine synthase subunit gamma |
| Hydrazine metabolism | 2151 | c_000000000027 | 28474 | 30886 | hzsA | Hydrazine synthase subunit alpha |
| Hydrazine metabolism | 2152 | c_000000000027 | 31134 | 32703 | hzo | Hydrazine dehydrogenase |

| | | | | | | |
|---|---|---|---|---|---|---|
| Hydrazine metabolism | 557 | c_000000000003 | 143276 | 144761 | hzo_2 | Hydrazine dehydrogenase |
| Hydrazine metabolism | 581 | c_000000000004 | 4711 | 5914 | hzo_3 | Hydrazine dehydrogenase |
| Hydroxylamine metabolism | 968 | c_000000000007 | 27484 | 29263 | hao1_1 | Hydroxylamine oxidoreductase |
| Hydroxylamine metabolism | 1046 | c_000000000008 | 3545 | 5246 | hao1_2 | Hydroxylamine oxidoreductase |
| Hydroxylamine metabolism | 2092 | c_000000000026 | 4158 | 5964 | hao1_3 | Hydroxylamine oxidoreductase |
| **Sulfur metabolism** | | | | | | |
| Sulfate Transport | 1378 | c_000000000012 | 30656 | 31367 | cysA | Sulfate/thiosulfate import ATP-binding protein CysA |
| Sulfate Reduction | 1376 | c_000000000012 | 28282 | 29596 | | Dissimilatory sulfite reductase |
| Sulfate Reduction | 2160 | c_000000000027 | 38291 | 40583 | phsA | Thiosulfate reductase molybdopterin-containing subunit PhsA |
| Sulfate Reduction | 2265 | c_000000000030 | 27976 | 29158 | sat | Sulfate adenylyltransferase |
| Sulfate Reduction | 2266 | c_000000000030 | 29160 | 29490 | aprB | Adenylylsulfate reductase subunit beta |
| Sulfate Reduction | 2267 | c_000000000030 | 29494 | 31177 | aprA | Adenylylsulfate reductase subunit alpha |
| Sulfate Reduction | 156 | c_000000000001 | 166210 | 167044 | asrB | Anaerobic sulfite reductase subunit B |
| Sulfate Reduction | 157 | c_000000000001 | 167046 | 168084 | asrA | Anaerobic sulfite reductase subunit A |
| Sulfate Reduction | 959 | c_000000000007 | 19766 | 20657 | asrC | Anaerobic sulfite reductase subunit C |
| Sulfate Reduction | 618 | c_000000000004 | 38750 | 39104 | dsrE | Putative sulfurtransferase DsrE |
| **Flagellar synthesis** | | | | | | |
| Flagellar synthesis | 1408 | c_000000000012 | 56831 | 57197 | flgD | Basal-body rod modification protein FlgD |
| Flagellar synthesis | 1409 | c_000000000012 | 57251 | 58862 | flgE | Flagellar hook protein FlgE |
| Flagellar synthesis | 1410 | c_000000000012 | 59180 | 59900 | flgG_1 | Flagellar basal-body rod protein FlgG |
| Flagellar synthesis | 1411 | c_000000000012 | 59926 | 60712 | flgG_2 | Flagellar basal-body rod protein FlgG |
| Flagellar synthesis | 1412 | c_000000000012 | 60761 | 61799 | flgA | Flagellar basal-body P-ring protein FlgA |
| Flagellar synthesis | 1413 | c_000000000012 | 61844 | 62504 | flgH | Flagellar L-ring protein |

| | | | | | | |
|---|---|---|---|---|---|---|
| Flagellar synthesis | 1414 | c_000000000012 | 62760 | 63885 | flgI | Flagellar P-ring protein |
| Flagellar synthesis | 1415 | c_000000000012 | 63931 | 64252 | flgJ | Peptidoglycan hydrolase FlgJ |
| Flagellar synthesis | 1416 | c_000000000012 | 64641 | 65208 | flgN | Flagellar protein FlgN |
| Flagellar synthesis | 1417 | c_000000000012 | 65580 | 67752 | flgK | Flagellar hook-associated protein 1 |
| Flagellar synthesis | 1418 | c_000000000012 | 67771 | 68674 | flgL | Flagellar hook-associated protein 3 |
| Flagellar synthesis | 1419 | c_000000000012 | 69017 | 69470 | fliW | Flagellar assembly factor FliW |
| **Carbon metabolism** | | | | | | |
| Glycogen Formation/Degradation | 716 | c_000000000004 | 139432 | 140581 | glgA | Glycogen synthase |
| Glycogen Formation/Degradation | 245 | c_000000000001 | 270426 | 271668 | glgC | Glucose-1-phosphate adenylyltransferase |
| Glycogen Formation/Degradation | 1058 | c_000000000008 | 18417 | 20004 | algC | Phosphomannomutase/phosphoglucomutase |
| Glycoxylate Shunt | 689 | c_000000000004 | 109210 | 109828 | gph_1 | Phosphoglycolate phosphatase |
| Glycoxylate Shunt | 833 | c_000000000005 | 124045 | 124738 | gph_2 | Phosphoglycolate phosphatase |
| Glycoxylate Shunt | 1641 | c_000000000016 | 35232 | 35919 | cbbZP | Phosphoglycolate phosphatase |
| Glycoxylate Shunt | 582 | c_000000000004 | 6030 | 6951 | mdh | Malate dehydrogenase |
| Pentose-phosphate pathway | 1319 | c_000000000011 | 41911 | 43441 | zwf2 | Glucose-6-phosphate 1-dehydrogenase 2 |
| Pentose-phosphate pathway | 1320 | c_000000000011 | 43498 | 44920 | gnd | 6-phosphogluconate dehydrogenase decarboxylating |
| Pentose-phosphate pathway | 1321 | c_000000000011 | 44989 | 45745 | pgl | 6-phosphogluconolactonase |
| Pentose-phosphate pathway | 1322 | c_000000000011 | 45758 | 47792 | tkt | Transketolase |
| Pentose-phosphate pathway | 1323 | c_000000000011 | 47845 | 48871 | glk | Glucokinase |

| | | | | | | |
|---|---|---|---|---|---|---|
| Glycolysis/Gluconegenesis | 1316 | c_000000000011 | 39873 | 40851 | pyk | Pyruvate kinase |
| Glycolysis/Gluconegenesis | 1058 | c_000000000008 | 18417 | 20004 | algC | Phosphomannomutase/phosphoglucomutase |
| Glycolysis/Gluconegenesis | 2084 | c_000000000025 | 43627 | 44908 | eno | Enolase |
| Glycolysis/Gluconegenesis | 1618 | c_000000000016 | 9729 | 10332 | hxlB | 3-hexulose-6-phosphate isomerase |
| Glycolysis/Gluconegenesis | 1620 | c_000000000016 | 10864 | 11581 | deoC1 | Deoxyribose-phosphate aldolase 1 |
| Glycolysis/Gluconegenesis | 1621 | c_000000000016 | 11573 | 12758 | deoB | Phosphopentomutase |
| Glycolysis/Gluconegenesis | 1696 | c_000000000017 | 30896 | 31670 | tpiA | Triosephosphate isomerase |
| Glycolysis/Gluconegenesis | 2048 | c_000000000025 | 667 | 2062 | pgi | Glucose-6-phosphate isomerase |
| Glycolysis/Gluconegenesis | 1323 | c_000000000011 | 47845 | 48871 | glk | Glucokinase |
| Glycolysis/Gluconegenesis | 1639 | c_000000000016 | 32672 | 34025 | glmM | Phosphoglucosamine mutase |
| Glycolysis/Gluconegenesis | 1640 | c_000000000016 | 34026 | 35214 | glmU_2 | Bifunctional protein GlmU |
| Glycolysis/Gluconegenesis | 1641 | c_000000000016 | 35232 | 35919 | cbbZP | Phosphoglycolate phosphatase |
| Glycolysis/Gluconegenesis | 1642 | c_000000000016 | 35951 | 37640 | | Gluconeogenesis factor |
| Glycolysis/Gluconegenesis | 433 | c_000000000003 | 11032 | 12220 | pgk_1 | Phosphoglycerate kinase |
| Glycolysis/Gluconegenesis | 434 | c_000000000003 | 12233 | 13235 | gap | Glyceraldehyde-3-phosphate dehydrogenase |
